# Change in RhoGAP and RhoGEF availability drives transitions in cortical patterning and excitability in *Drosophila*

**DOI:** 10.1101/2023.11.06.565883

**Authors:** Jonathan A. Jackson, Marlis Denk-Lobnig, Katherine A. Kitzinger, Adam C. Martin

**Affiliations:** Department of Biology, Massachusetts Institute of Technology; Graduate Program in Biophysics, Harvard University; Department of Biophysics, University of Michigan

**Keywords:** RhoA, actomyosin cortex, excitability, development, *Drosophila* egg chamber, *Drosophila* embryo

## Abstract

Actin cortex patterning and dynamics are critical for cell shape changes. These dynamics undergo transitions during development, often accompanying changes in collective cell behavior. While mechanisms have been established for individual cells’ dynamic behaviors, mechanisms and specific molecules that result in developmental transitions in vivo are still poorly understood. Here, we took advantage of two developmental systems in *Drosophila melanogaster* to identify conditions that altered cortical patterning and dynamics. We identified a RhoGEF and RhoGAP pair whose relocalization from nucleus to cortex results in actomyosin waves in egg chambers. Furthermore, we found that overexpression of a different RhoGEF and RhoGAP pair resulted in actomyosin waves in the early embryo, during which RhoA activation precedes actomyosin assembly and RhoGAP recruitment by ∼4 seconds. Overall, we showed a mechanism involved in inducing actomyosin waves that is essential for oocyte development and is general to other cell types.

## Introduction

Actomyosin contractility plays a key role in generating force during cell shape change and tissue morphogenesis. Proper shape requires that patterns of forces are tailored to the specific developmental context, and the forces required can even change within the same tissue at different stages of its morphogenesis^1–3^. A key regulator controlling these specific patterns of forces is the small GTPase RhoA. RhoA promotes actomyosin contractility by activating the Rho-associated protein kinase (ROCK), which activates non-muscle myosin II, and the Diaphanous family of formins, which induce unbranched actin assembly. RhoA can cycle between an active GTP-bound state and inactive GDP-bound state. This cycling is controlled by a battery of Rho guanine nucleotide exchange factors (RhoGEFs) and GTPase activating proteins (RhoGAPs), which promote GTP loading and hydrolysis, respectively (Figure 1A)^4^, often with several GEFs and GAPs existing in the same cell with different spatial and temporal behavior^5,6^. These RhoGEFs and RhoGAPs are themselves developmentally regulated and function in modular and combinatorial ways to spatially and temporally control actomyosin distribution^7^.

**Figure 1:**
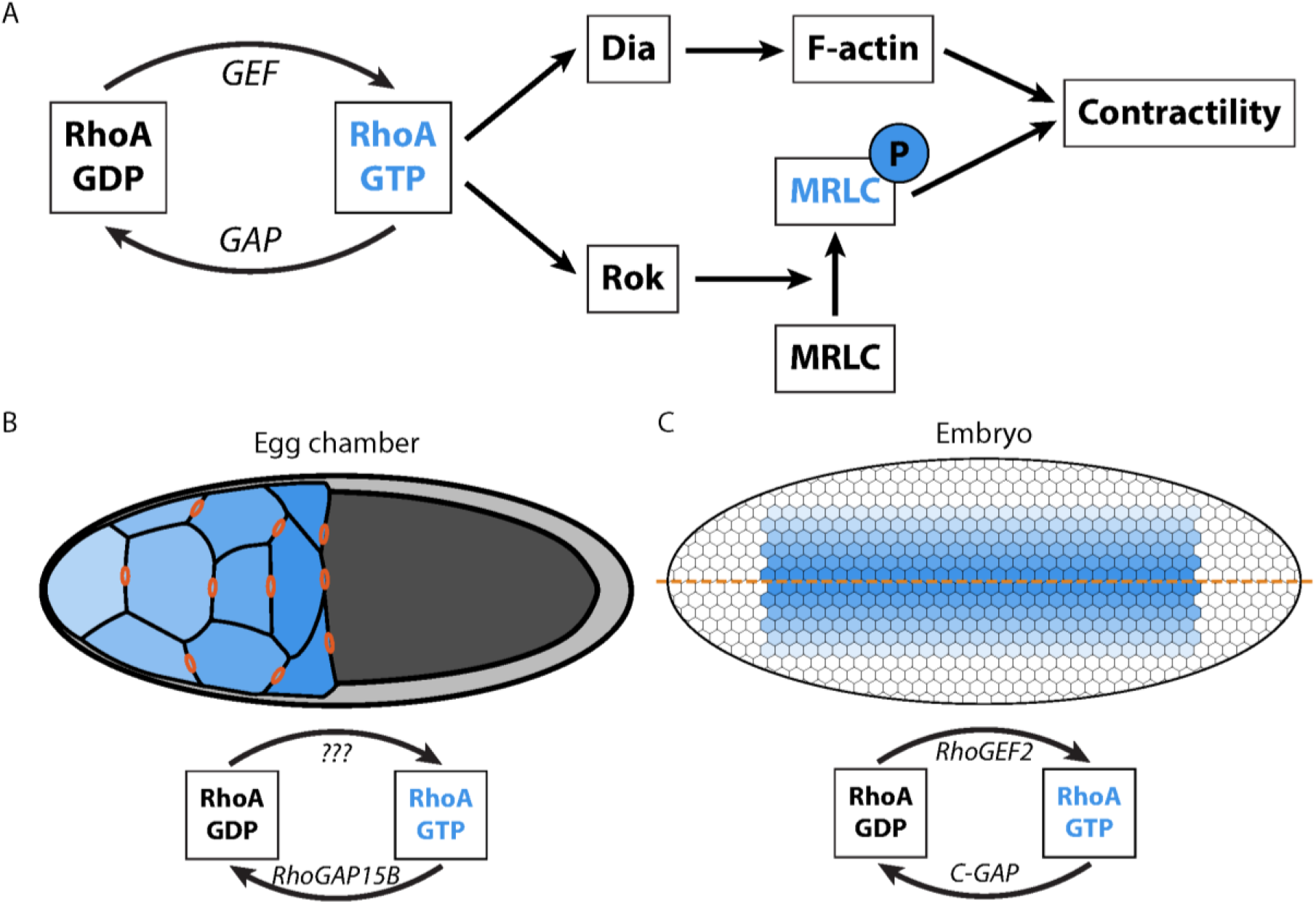
The RhoA pathway regulates actomyosin contractility in embryos and egg chambers. **A**. Subset of the components and interactions in the RhoA pathway in *Drosophila*. When GTP-bound, RhoA activates both Rho kinase (Rok) and the formin Diaphanous (Dia), which lead to myosin regulatory light chain (MRLC) phosphorylation, F-actin assembly, and actomyosin contractility. GEFs and GAPs activate and inactivate RhoA, respectively, and are themselves developmentally regulated. **B**. In the egg chamber (shown with anterior at left and ring canals in orange), actomyosin waves appear in nurse cells (blue) starting from the most posterior cells and appearing later in the more anterior ones. RhoGAP15B plays the role of the GAP; no GEF has so far been identified for these waves. **C**. Actomyosin pulses appear in the embryo in the rows of cells surrounding the ventral midline (orange line), with activity higher and appearing earlier near the midline^28^. RhoGEF2 and C-GAP form a regulatory pair for RhoA in this system. These cells are 5-10 µm in diameter, significantly smaller than the nurse cells, which are 30-40 µm in diameter when waves appear. For **B** and **C**, blue color represents the gradient of actomyosin appearance, where darker color implies earlier appearance.

The RhoA signaling system has been shown to exhibit feedback and excitability, including oscillatory and wave-like behaviors. Evidence suggests that RhoA can promote its own activation, creating an autocatalytic positive feedback loop that will drive the system to peak activity after crossing a threshold. Possible mechanisms for this positive feedback include cortical advection and/or contraction concentrating RhoA and its effectors^8,9^, and contraction-independent recruitment or activation of RhoA^10–12^. Delayed negative feedback terminates the RhoA signal, often requiring the activity of a RhoA GAP^10,13–17^. The timing or spatial position of this termination creates a refractory period or ‘dead’ zone where RhoA cannot be easily reactivated. Such zones lead to hallmarks of excitability, such as annihilation of waves upon collision. While the theory of excitability is well established and some examples of the signaling circuits that lead to excitability have been defined^8,14,15,18^, the specific sets of molecules that drive events in development and how the regulatory principles compare across cell types are still poorly understood.

There are developmental events during which there is a clear transition to excitable actomyosin behavior. In the *Drosophila* female germline, nurse cells that lack apical-basal polarity transition from having a stable actomyosin cortex to exhibiting traveling actomyosin waves. The germline develops as a cyst of interconnected cells with the nurse cells connected to the oocyte through actin-rich bridges called ring canals (Figure 1B), through which cytoplasm is transported to the oocyte in a process known as nurse cell dumping (hereafter referred to as ‘dumping’). In this system, actomyosin waves emerge after the nurse cells transfer most of their volume and are required to enable the transport of remaining cytoplasm to the oocyte. We previously identified a RhoGAP, RhoGAP15B, that is involved in regulation of wave behavior^13^, but the identity of a cognate RhoGEF that functions in this process was unknown. Nurse cells display a size hierarchy and stereotyped dumping onset order, with cell size decreasing and dumping proceeding later from posterior (nearer the oocyte) to anterior (farther from the oocyte)^13,19^. Actomyosin waves have been shown to appear in the same order as cell size decrease (Figure 1B, blue), but it is unknown what triggers the actomyosin waves and whether they emerge through a switch-like or gradual transition.

Dynamic actomyosin also plays a role in epithelial cells. During *Drosophila* gastrulation, presumptive mesoderm cells exhibit apical constriction, during which actomyosin contractility is targeted to the middle of the apical cell surface (medioapical)^20,21^. Mesoderm cells do not exhibit actomyosin waves, but stationary pulses of actomyosin assembly and disassembly. This is in contrast to other apically constricting cells that exhibit actomyosin waves across the apex^22,23^. Similar to the *Drosophila* germline, mesoderm cells exhibit a spatial pattern of actomyosin across the tissue, with actomyosin assembly initiating earlier and with higher levels along the ventral midline and with more dynamic behavior in more lateral mesoderm cells^24–27^ (Figure 1C). Actomyosin pulses appear to be activated by RhoGEF2, a RhoGEF that is specific to RhoA. A RhoGAP, Cumberland GAP (C-GAP, also known as RhoGAP71E), terminates RhoA signaling to disassemble the actomyosin patch^14^. Whether these different dynamics between cells reflect differences between the specific GEFs and GAPs involved, epithelial vs. non-epithelial cell type, distinct RhoA activity, or GEF and GAP activity is unclear.

Here, we investigated the regulatory systems that are used to alter actomyosin state in the *Drosophila* germline and early embryo. We identify a RhoGEF, Pebble (the *Drosophila* Epithelial cell transforming 2 (Ect2) homologue), involved in nurse cell actomyosin wave formation and show that actomyosin wave induction is associated with Ect2/Pebble and RhoGAP15B nuclear release. We show that co-overexpressing a different RhoGEF and RhoGAP pair, RhoGEF2 and C-GAP, can induce actomyosin waves in epithelial cells and the early embryo, suggesting that such excitable behavior does not depend on specific RhoGEF and RhoGAP proteins or cell size/type and instead might depend on the structure of the RhoA regulatory network. Furthermore, analysis of actomyosin waves in the embryo supports similar regulatory logic for RhoGEF2 and C-GAP induced waves as that shown for actomyosin waves in *C. elegans*, *Xenopus*, and starfish, suggesting a common regulatory logic for the RhoA system across diverse systems, cell types, and molecular identities of RhoGEFs and RhoGAPs.

## Results

### Wild-type Ect2 expression is required for actomyosin waves in nurse cells

To investigate mechanisms that promote the transition from uniform contractility to actomyosin waves in nurse cells, we sought to identify RhoGEFs involved in dumping. First, we assessed embryo size upon germline-specific depletion of individual RhoGEFs by RNA interference (see Methods for details), as incomplete dumping can lead to smaller embryos^13,29,30^. We previously used RNAi screening to identify a RhoGAP (RhoGAP15B) required for dumping^13^, which we included in the screen as a positive control along with the adducin homologue *hu li tai sho* (*hts*)^30^. Short-embryo phenotypes were rare even for positive controls, presumably because knockdowns were incomplete. Of the RhoGEF lines that occasionally resulted in short embryos (Figure 2A-B), we chose to focus on Ect2 because both Ect2-RNAi lines resulted in a higher phenotypic frequency than our negative control, Ect2 has been shown to induce actomyosin waves and pulses in other systems^16,31,32^, and Ect2’s localization suggested its function in dumping (see below). Ect2 is most commonly associated with the regulation of RhoA in cytokinesis^33^; however, we confirmed that knockdowns affected dumping by live imaging using a GFP-tagged version of the regulatory light chain of non-muscle myosin II (Spaghetti squash, Sqh). Of the candidate RhoGEFs in Figure 2A, knockdown of Ect2 most frequently led to erratic or blocked waves and to a failure of dumping similar to that seen upon knockdown of RhoGAP15B^13^ (Figure 2C-E; Supplemental Figure S1A-F; Video S1).

**Figure 2:**
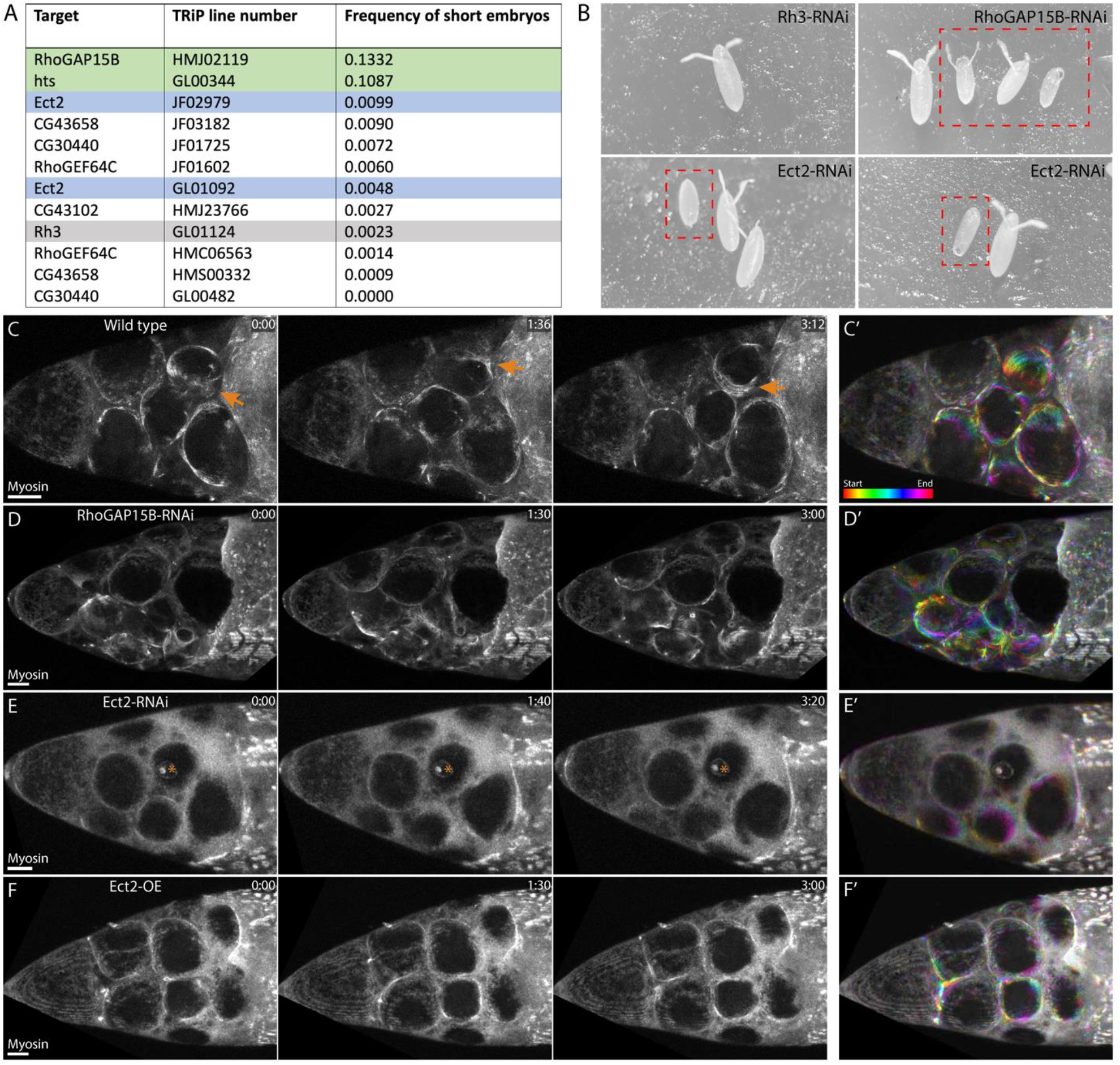
Ect2 and RhoGAP15B are required for proper regulation of actomyosin waves. **A**. Results from a RhoGEF RNAi screen in fly lines for which short embryos were occasionally observed. Green rows are positive controls; gray row is a negative control. Two different Ect2 lines were screened, with slightly different frequencies of short embryos. Egg chambers from lines with short-embryo frequencies above that of the negative control were also imaged live to determine if knockdown affected waves. **B**. Examples of short embryos (red boxes) next to normal-length embryos for negative control (Rh3-RNAi), a positive control (RhoGAP15B-RNAi), and two separate examples of Ect2-RNAi. **C**. Maximum-intensity projections (MIPs) of myosin signal from a movie of a wild-type egg chamber in which multiple cells display myosin waves. Orange arrows point to a wave propagating across the cortex of one cell. **D**. MIPs from a RhoGAP15B-RNAi egg chamber. Dumping begins normally, but stalls as a subset of cells becomes smaller than others, and several cells display erratic waves (cyan arrows) that do not propagate fully across the cortex. **E**. MIPs from an Ect2-RNAi egg chamber. Cells round up slightly as usual preceding actomyosin wave appearance, but few myosin accumulations appear and dumping stalls. Asterisks mark myosin signal from stalk cells, which are external to the germline. **F**. MIPs from an Ect2-OE egg chamber. Similar to RhoGAP15B-RNAi, dumping stalls as some cells develop small, erratic myosin accumulations (cyan arrows). Scale bars for **C**-**F**: 20 µm. Time stamps are in min:sec. **C’**-**F’** show color-coded projections of the same egg chambers as in C-F through 2 minutes, starting at the earliest time point shown. Color bar in **C’** indicates order of frames. All MIPs are through slightly over half of the egg chamber, and in all images, the oocyte, not shown, is to the right.

**Supplemental Figure S1:**
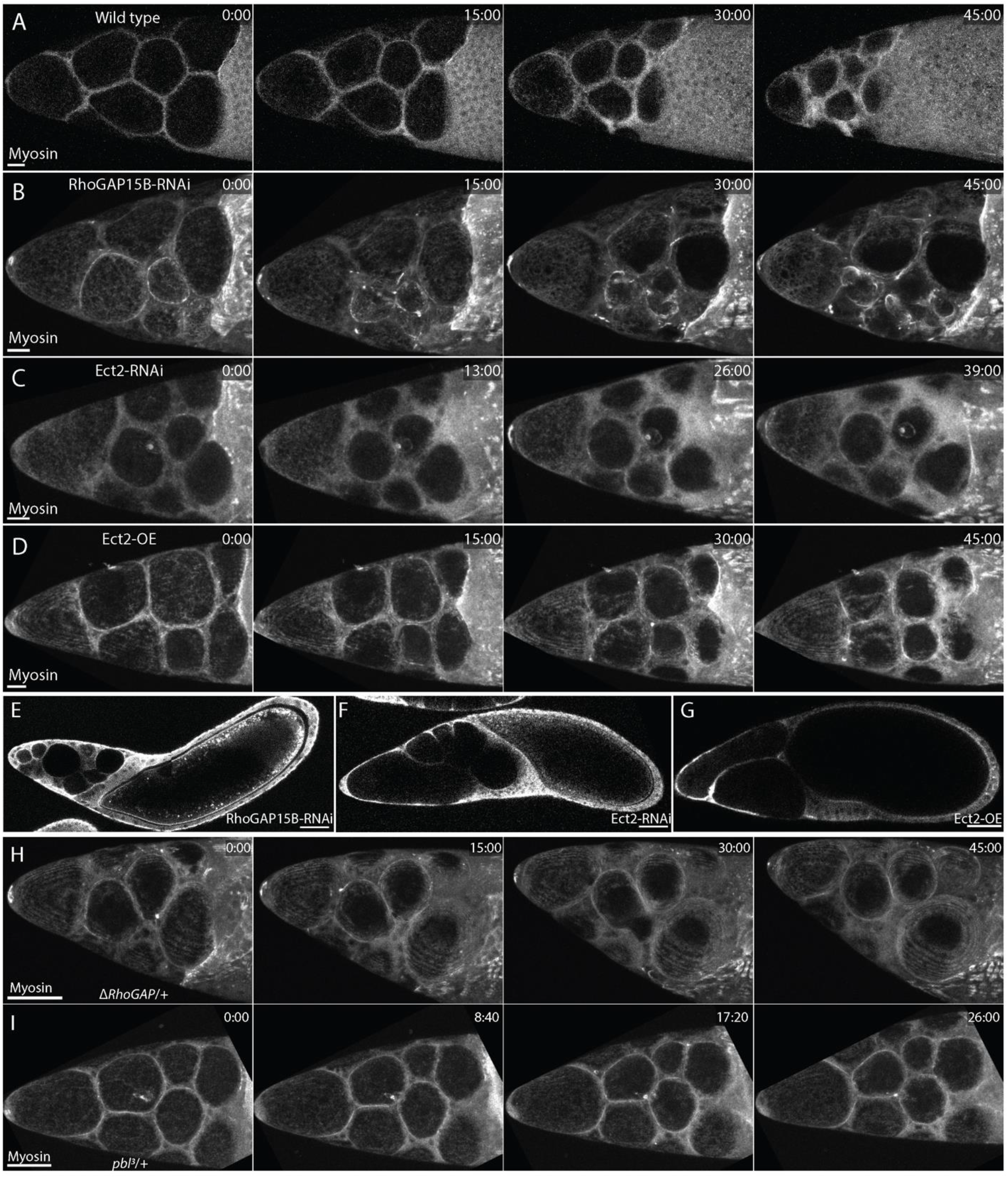
Ect2 and RhoGAP15B perturbations block nurse cell dumping. **A**. MIPs of myosin signal in a wild-type egg chamber showing reduction in cell volume and appearance of nonuniform myosin intensity in the third and fourth time points. **B**. MIPs from the same egg chamber as in Figure 1D, showing dumping stalling as erratic waves form in the smaller cells in the egg chamber. **C**. MIPs from the same egg chamber as in Figure 1E, showing dumping stalling without large-scale wave formation. **D**. MIPs from the same egg chamber as in Figure 1F. Note the reduction in nurse cell cluster size in **A** that is not seen to the same degree in **B**-**D**. **E**-**G**. Images of egg chambers after dumping has stalled for (**E**) RhoGAP15B-RNAi, (**F**) Ect2-RNAi, and (**G**) Ect2-OE egg chambers. **E** and **F** are single optical sections from near the midplane, while **G** is a maximum-intensity projection. **H**, **I**. MIPs of myosin signal in egg chambers heterozygous for an allele of *RhoGAP15B* missing its RhoGAP domain (abbreviated as *ΔRhoGAP*) or for *pbl*^3^, a null allele of Ect2. Scale bars: 20 µm in **A**-**D**; 50 µm in **E**-**I**. Time stamps are min:sec.

To further support the role of RhoGAP15B and Ect2 as a GAP/GEF pair regulating actomyosin waves, we depleted each gene using egg chambers heterozygous for an allele of RhoGAP15B in which the RhoGAP domain was deleted via CRISPR (see Methods) or egg chambers heterozygous for *pbl*^3^, a null allele of *Ect2*^34^. We chose to focus on heterozygous rather than homozygous mutants both because homozygous *pbl*^3^ is lethal and we expected heterozygous mutants to reduce gene dosage in a way that would be comparable to RNAi. In both heterozygous mutants, waves appeared less continuous than in wild-type chambers, and waves often did not appear at all in *pbl*^3^ (Supplemental Figure S1H,I; Video S1). Additionally, overexpression of Ect2 in the germline (Ect2-OE) resulted in nurse cell waves that were shorter-lived than in wild type and frequently led to incomplete dumping (Figure 2F; Supplemental Figure S1G; Video S1), suggesting that the levels and/or regulation of Ect2 dynamics are important for the dumping process.

Prior work has shown that nurse cell dumping proceeds in two phases: in the first, transport is driven by cell size and pressure differences, with myosin roughly uniform along the cortex, while the second phase requires actomyosin waves to complete dumping^13^. In all knockdowns studied here, cytoplasmic transport occurs at least to some extent, consistent with this two-phase model of dumping. RhoGAP15B reduction does not obviously perturb dumping until the first cells begin to develop actomyosin contractions, after which cells often fuse and dumping is blocked^13^ (Figure 2D; Supplemental Figure S1H; Video S1). Nurse cells with reduced Ect2 levels transfer some of their contents in the first phase of dumping but frequently fail to develop waves and transition to the second phase (Figure 2E; Supplemental Figure S1I; Video S1), consistent with the two-phase model and suggesting Ect2 is important for wave onset. Stalled cytoplasmic transfer upon RhoGAP15B or Ect2 perturbation is unlikely to be the result of blocked ring canals. Although ring canals connecting nurse cells and the oocyte are formed from stalled cytokinetic furrows, and Ect2 plays a role in furrow constriction^35,36^, ring canals did not appear obstructed or different from wild-type canals in either RhoGAP15B-RNAi^13^ or *pbl*^3^ heterozygous mutants (not shown).

Ect2, possibly alongside other GEFs, is thus involved in the regulation of actomyosin waves in the nurse cells. Together with our previous identification of RhoGAP15B and our analysis of a new RhoGAP15B allele here, our results suggest that both Ect2 and RhoGAP15B regulate actomyosin wave formation. Because actomyosin reorganizes during nurse cell dumping, we next examined whether the localization of these two proteins changes with the onset of actomyosin waves.

### Nuclear exit of RhoGEF/Ect2 and RhoGAP15B is associated with actomyosin wave onset

To determine whether Ect2 and RhoGAP15B levels or localization change when the actomyosin cortex transitions from uniform to wave-like behavior, we imaged labeled versions of both proteins during the dumping process. Prior to wave onset, Ect2 was localized to the nucleus in nurse cells, but exited the nucleus and became enriched at the cortex as dumping proceeded (Figure 3A,A’; Video S2). RhoGAP15B showed the same transition from nucleus to cortex: although there was cortical enrichment prior to dumping onset, the nuclear signal decreased significantly while the intensity increased at the cortex with time (Figure 3B,B’; Video S2). To quantify these observations, we measured the ratio of cortical to nuclear intensity of each protein around dumping onset (‘early’) and near wave onset (‘late’), finding that the intensity ratio increased approximately 9-fold for Ect2 and 3-fold for RhoGAP15B (Figure 3C).

**Figure 3:**
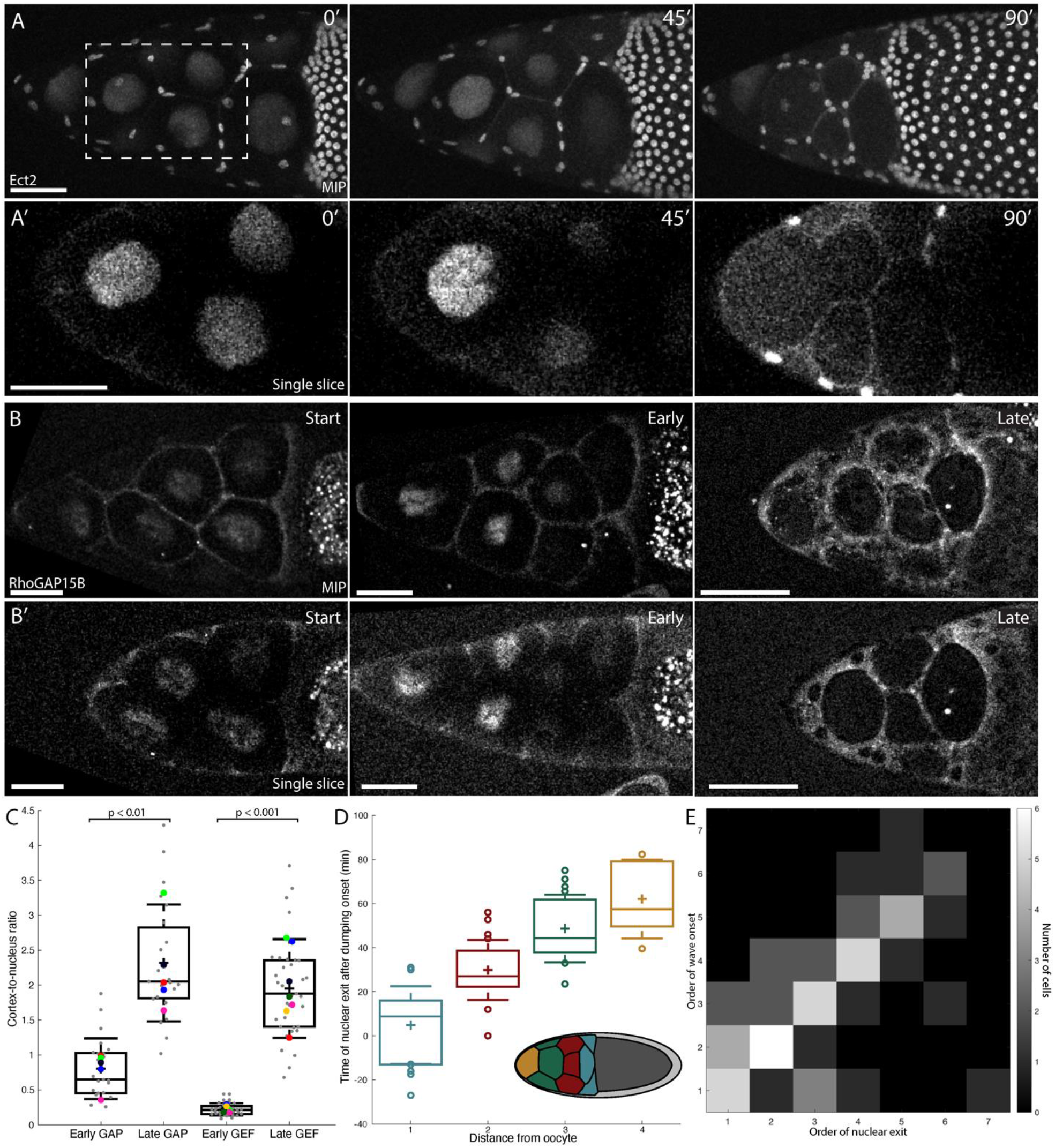
Ect2 and RhoGAP15B exit the nucleus prior to and in the same order as wave appearance. **A**. MIPs of Ect2::EGFP signal in an egg chamber at three time points in dumping, with corresponding zoom-ins of a single optical section from the boxed region in **A’**. **B**. MIPs of RhoGAP15B::sfGFP signal from three separate egg chambers just after the start of dumping, early in dumping, and late in dumping, roughly analogous to the three time points in **A** based on size and shape of the nurse cell cluster. Single optical sections from each egg chamber are shown in **B’**. **C**. Ratio of cortical to nuclear intensity of RhoGAP15B and Ect2 early and late in dumping. Sample sizes: 23 cells from 5 egg chambers (RhoGAP15B) and 35 cells from 7 egg chambers (Ect2). p-values are 0.008 and 6 x 10^−4^ for GAP and GEF, respectively (Mann-Whitney U test). **D**. Time after dumping onset at which PCNA becomes noticeably cytoplasmic (details in Methods), grouped by number of ring canals separating a nurse cell from the oocyte. As with wave order, cells nearer the oocyte show earlier leakage of contents from the nucleus. Inset: schematic of egg chamber with groups of nurse cells colored by distance from the oocyte, corresponding to boxplot colors. **E**. Heat map of order of PCNA exit from the nucleus versus order of wave onset. Data for **D**,**E** come from 11 egg chambers, with 14, 20, 16, and 5 cells for distances 1, 2, 3, and 4 from the oocyte, respectively. All scale bars: 50 µm. For box plots, center line denotes median, plus sign mean, box edges 25th and 75th percentile, and whiskers standard deviation. In C, colored dots show mean ratio per egg chamber; gray points are individual cells. In **D**, only individual points outside mean +/− standard deviation are shown.

**Supplemental Figure S2:**
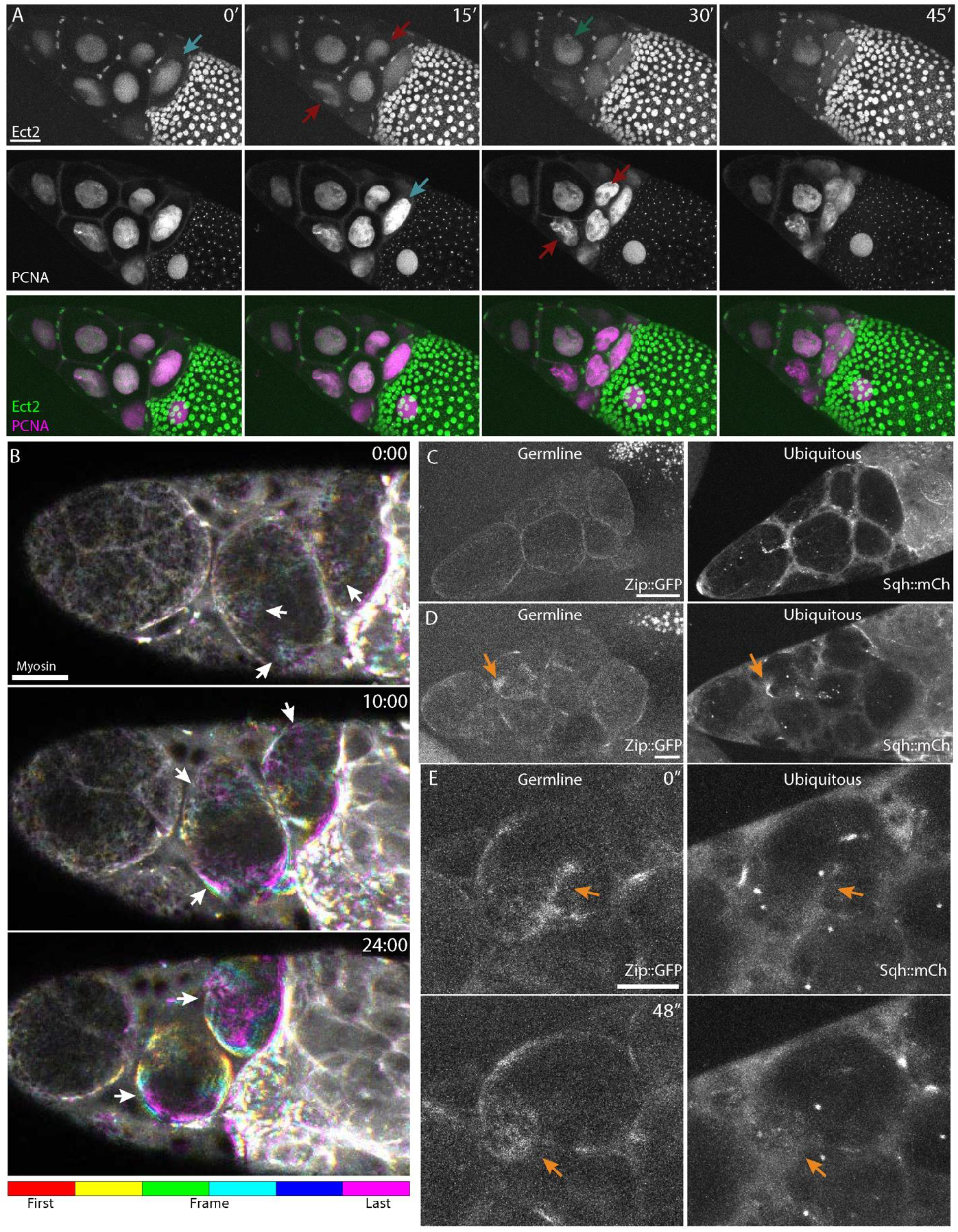
PCNA serves as a proxy for Ect2 release from the nucleus as myosin gradually becomes more wave-like. **A**. MIPs of Ect2 (top row) and PCNA (middle row) during dumping for one representative egg chamber, with merged image in the bottom row. Colored arrows point to approximate exit from the nucleus, with blue, red, or green arrows corresponding to cells one, two, or three ring canals from the oocyte, respectively. PCNA exits the nucleus roughly 10-15 minutes after but in the same order as Ect2. **B**. Three MIPs of myosin signal in an egg chamber with six subsequent frames, spanning one minute total, shown in different colors. From an initially uniform cortical distribution, myosin first shows temporary spatial accumulations or ‘flickers’ (top panel, arrows). Later in dumping, the flickers develop into faint waves (middle panel, arrows) that become progressively more persistent and intense (bottom panel, arrows). **C**. Germline-specific myosin signal (myosin heavy chain, Zipper::GFP; left) and ubiquitously-expressed myosin signal (Sqh::mCherry; right) in an egg chamber just prior to wave onset. **D**. Images from the same egg chamber following wave onset. **E**. Zoom-in on the cell highlighted by the arrow in **D**. Arrows in **D**,**E** point to waves. Scale bars: 50 µm (**A**-**C**) and 20 µm (**D**,**E**).

To determine if the timing of Ect2 nuclear exit correlates with the timing of wave onset, we asked whether exit of Ect2 from the nucleus in different cells might follow the same posterior-to-anterior order as wave onset. Slight Ect2 overexpression sometimes affected wave behavior and led to cell fusion, complicating quantitative analysis. However, PCNA, a sliding clamp for DNA polymerase, normally localizes to the nucleus in interphase cells^37^ but has been shown to exit the nucleus in the same order as dumping onset^13^. Because PCNA is not expected to interfere with the RhoA pathway, it serves as an ideal proxy for RhoGAP15B and Ect2 exit from the nucleus. Indeed, PCNA exits nuclei in the same order as Ect2 but does so 10-15 minutes later (Supplemental Figure S2A). Grouping PCNA exit time by the number of cytoplasmic bridges between nurse cells and the oocyte revealed the same order as wave onset, in which cells nearest the oocyte show leakage of contents from the nucleus first (Figure 3D). Further, a heatmap of the ranked order in which PCNA exits the nucleus and the order in which waves begin in nurse cells shows that rank pairs are distributed around a diagonal line (Figure 3E). Both findings suggest exit of proteins from the nucleus and wave onset are correlated in the order they occur at the level of individual nurse cells, consistent with the hypothesis that relocalization of GEF and GAP from the nucleus to the cortex could trigger wave-like actomyosin behavior.

To test whether the onset of myosin wave dynamics might reflect the gradual release of Ect2 and RhoGAP15B from the nucleus, we examined nurse cells at higher spatial and temporal resolution to determine whether the onset of waves is gradual or switch-like. Myosin signal starts as a relatively static, reticular pattern before small flickers of brighter intensity appear apparently randomly over the network. As dumping progresses, faint bands of myosin appear from these flickers, subsequently becoming larger, more intense, and associated with greater cell deformation over time (Supplemental Figure S2B; Video S3). The Sqh::GFP construct used here expresses tagged myosin in all cell types, including the overlying somatic follicle cells that surround the nurse cell cluster. However, although these cells display their own myosin dynamics, we confirmed the flickers and waves studied here are present in the nurse cells using germline-specific Gal4 to express UAS-driven, GFP-tagged Zipper (myosin heavy chain; Supplemental Figure S2C-E, Video S4). In contrast, Zipper::GFP expressed specifically in the follicle cells showed neither waves nor the network-like myosin pattern seen in nurse cells prior to wave onset (not shown). Overall, the gradual onset of actomyosin waves specifically in the nurse cells supports a model in which GAP/GEF release induces a gradual transition to and strengthening of waves.

### Manipulation of a different RhoGEF and RhoGAP pair in the early embryo enables titration of RhoA activity

The importance of Ect2 and RhoGAP15B function in actomyosin wave propagation, as well as the correlation of Ect2 and RhoGAP15B cortical relocalization to the timing and progression of wave onset, suggested that increased cortical availability of these proteins is responsible for the transition in actomyosin behavior in nurse cells. To more broadly investigate how changing availability of GEF and GAP affects the mode of actomyosin behavior and whether this phenomenon is common to a broader set of GEFs, GAPs, and cell types, we next examined the effects of changing GEF and GAP levels in the early embryo around the time of gastrulation.

We have previously shown that we can modulate RhoA activity by modulating the expression of RhoGEF2 or C-GAP^14,24^. Therefore, we sought to calibrate the RhoA signaling level of mesoderm cells for various expression levels of GEF and GAP. We used medioapical myosin in mesoderm cells as a proxy for RhoA activity level, because its presence is dependent on RhoA activity^14^, medioapical myosin disappears within tens of seconds of Rok inhibition^38^, and myosin intensity is correlated with apical contractility^39^. The fraction of mesoderm cell apical area covered by myosin was correlated with the size and intensity of a GFP-tagged anillin Rho-binding domain^8^ (anillinRBD), consistent with myosin spot size reflecting RhoA activity (Supplemental Figure 3A-D). We then measured the fraction of the mesoderm cells’ apical surface covered by myosin spots by segmenting the myosin signal in projections of these cells. We found that overexpressing C-GAP or knocking down RhoGEF2 reduced coverage fraction, consistent with reduced RhoA activity (Video S4). In contrast, overexpressing RhoGEF2 resulted in larger and more prominent coverage of myosin spots across the tissue, consistent with increased RhoA activity (Figure 4A,B and Supplementary Figure 3A-C; Video S4). Knocking down C-GAP appeared to increase the amount of the cell surface covered by myosin (Figure 4A), although this change disrupted spot formation and was not reflected in our quantification (Figure 4B). This discrepancy may result from the more diffuse myosin signal upon C-GAP knockdown^14^ being omitted by our segmentation workflow (see Methods). While imaging RhoGEF2-overexpression (‘RhoGEF2-OE’) embryos, we noticed differences between the two existing fly lines (referred to here as RG2-OE(5) and RG2-OE(6b)) carrying the same overexpression construct on different chromosomes; of the two, RG2-OE(6b) had more severe effects (see Methods for details). For quantification of myosin coverage, only RG2-OE(5) embryos were used. Finally, we observed that co-overexpressing RhoGEF2 and C-GAP (‘double-OE’) suppressed the spot size changes of either overexpression alone (Figure 4B,C). Thus, individually modulating GEF and GAP expression levels leads to a continuum of different RhoA activity levels as expected based on the known function of these genes as activator and inhibitor, respectively.

**Figure 4:**
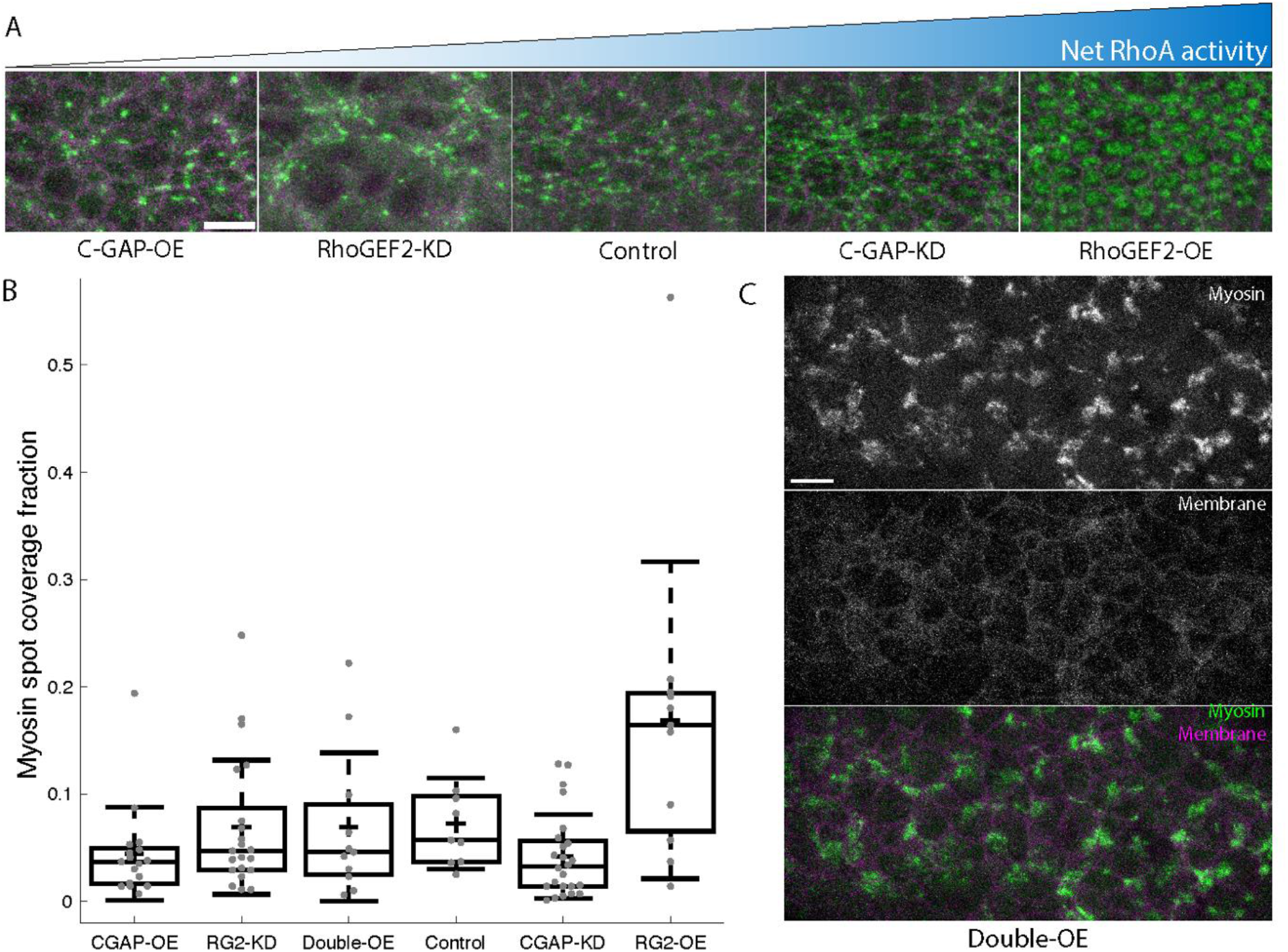
Myosin area coverage in mesoderm cells scales with net RhoA activity. **A**. Representative MIPs of myosin signal approximately 7.5 minutes after pulse appearance in five genotypes. **B**. Fraction of the mesoderm cells’ apical surface covered by myosin spots ∼7.5 minutes after pulse appearance. From left to right, n = 17, 21, 11, 9, 24, and 11 embryos. **C**. Myosin (top), membrane (middle), and merged (bottom) signal from an embryo overexpressing RhoGEF2 and C-GAP. All scale bars: 10 µm. KD = knockdown; OE = overexpression. In all images, myosin is in green and membranes are in magenta.

**Supplemental Figure S3:**
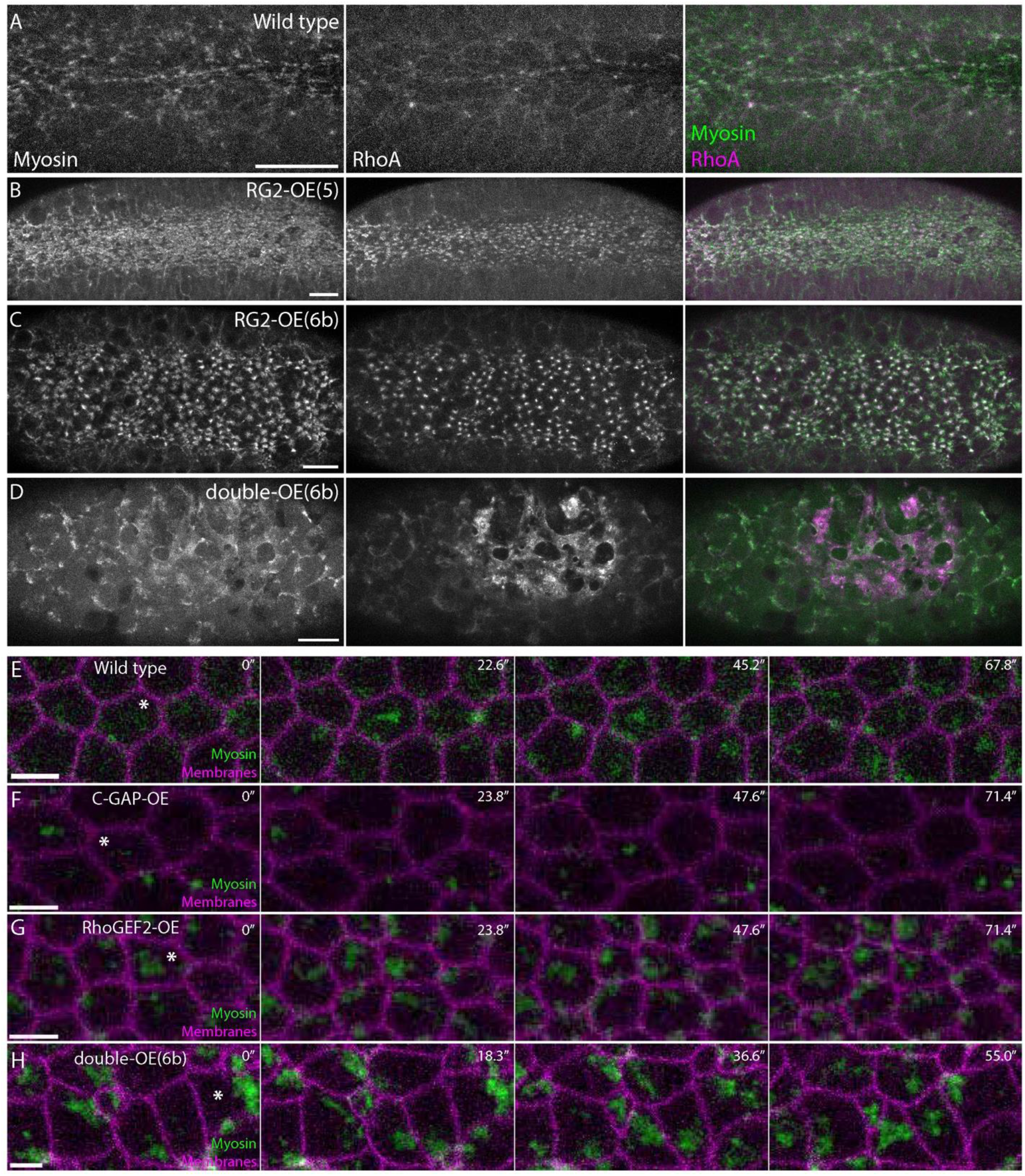
RhoA and myosin change upon perturbation to RhoGEF2 and C-GAP levels. **A**. MIPs of myosin (left), RhoA (center, visualized by the Rho-binding domain of anillin), and merged channels (right) in a wild-type embryo. Both proteins appear as spots within a supracellular network. **B**. Upon RhoGEF2 overexpression, both proteins form larger spots. **C.** Reorganization into spots is also seen in RhoGEF2-OE embryos in which gastrulation defects are more severe. **D.** Upon overexpression of both regulators, myosin and RhoA reorganize into traveling waves rather than spots. The bright region in the RhoA signal is the residual cellularization front. **E**. Time series from a wild-type embryo showing myosin pulsing and becoming more prominent over time. **F**. Myosin pulses in C-GAP-OE embryos are smaller and build up less over time. **G**. Myosin pulses in RhoGEF2-OE embryos are larger but still accumulate over time. **H**. Myosin in double-OE embryos is organized in traveling waves instead of medioapical pulses. Asterisks in **E**-**H** highlight cells with prominent examples of pulses or waves. Images in **E**-**H** are surface projections of the myosin channel and a single optical section of the membrane channel. Scale bars: 20 µm (**A**-**D**); 5 µm (**E**-**H**).

### Co-overexpression of RhoGEF2 and C-GAP leads to robust actomyosin wave formation

If RhoA activity and myosin behavior are determined by the ratio of GEF to GAP (analogous to the net RhoA activity), overexpressing RhoGEF2 in a C-GAP-OE background would suppress the C-GAP-OE phenotype. While co-overexpression did suppress myosin spot size changes (Figure 4B,C), we found that co-overexpressing RhoGEF2 and C-GAP in embryos (hereafter termed ‘double-OE embryos’) resulted in traveling myosin and RhoA waves both apically and laterally in mesoderm cells (Figure 5A; Supplemental Figure 3D-H; Video S5). Actomyosin waves did not promote sustained apical constriction, but instead transiently deformed cells. Thus, co-overexpression of RhoGEF2 and C-GAP, despite resulting in intermediate RhoA activity based on myosin spot size, resulted in a unique actomyosin behavior for mesoderm cells.

**Figure 5:**
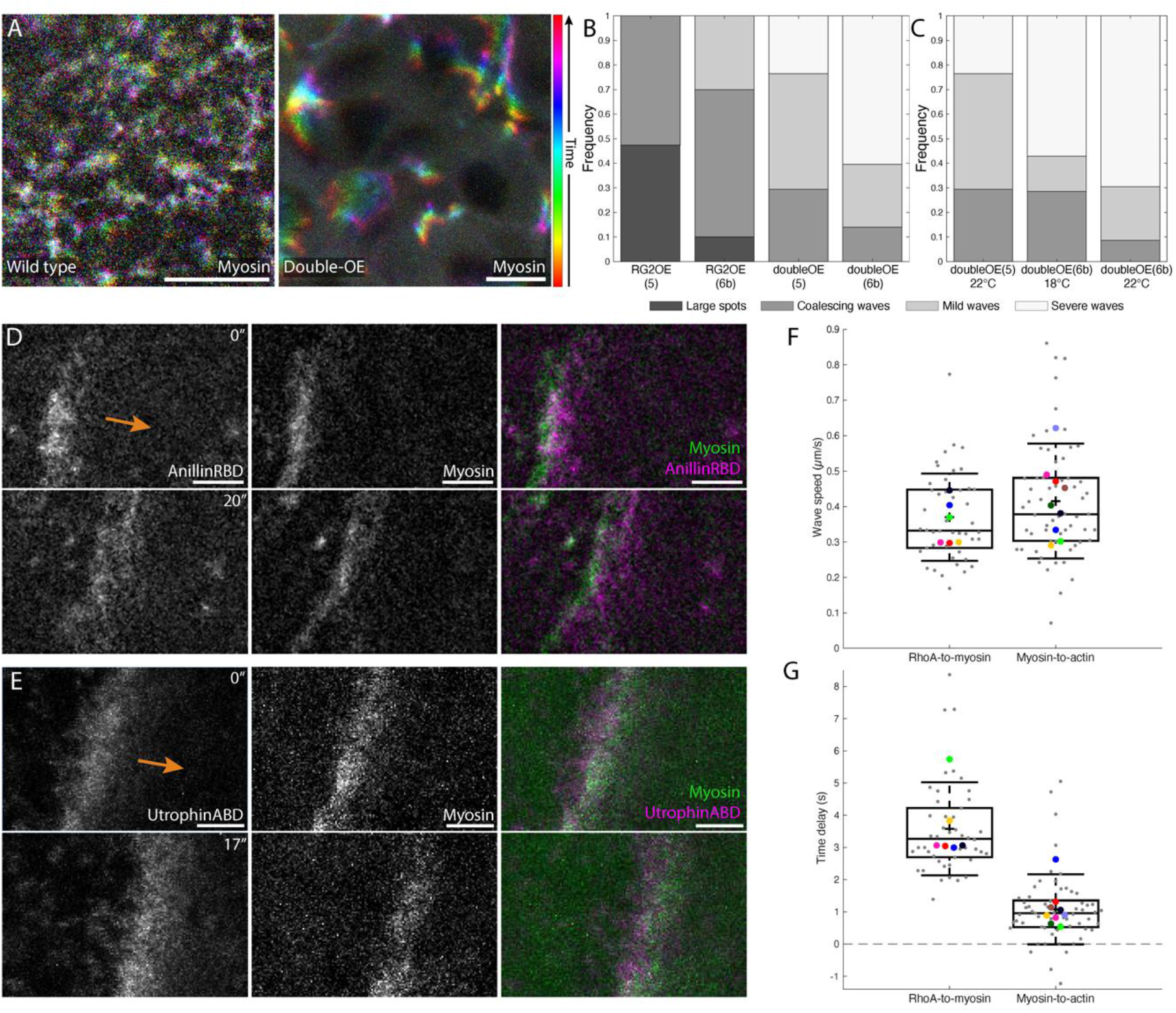
Wave phenotype severity scales with GAP and GEF availability, with severe waves revealing the order of RhoA pathway components. **A**. Projections of myosin signal for a wild-type embryo (left) and double-OE embryo (right), colored by time, spanning about 30 seconds. **B**. Frequency of different classes of phenotypes for RhoGEF2-OE and double-OE embryos (see Supplemental Figure 4 for details). n = 19, 10, 17, and 43 embryos for genotypes from left to right. **C**. Subset of data from **B**, with frequency of different classes of phenotypes for double-OE embryos grouped by temperature; overexpression strength is expected to increase with higher temperature^40^. From left to right, n = 17, 14, and 23 embryos. **D**. RhoA (left; visualized by GFP::AnillinRBD^8^), myosin (middle), and merged image (right) at two timepoints from an embryo following cellularization failure. RhoA leads myosin slightly in space, likely reflecting its earlier recruitment. Arrow indicates wave direction. **E**. F-actin (left; visualized using UtrABD::GFP), myosin (center), and merged image (right) from a different embryo, in which myosin slightly leads F-actin. **F**. Wave speed in embryos after cellularization failure. Difference in wave speed between the two conditions is not significant (p = 0.22, Mann-Whitney U test). **G**. Delay time between pairs of components from the same embryos as in **F**. Components in box plots in **F** and **G** are the same as for the box plots in Figure 3C; n = 46 waves from 6 embryos for RhoA-to-myosin and 61 waves from 9 embryos for actin-to-myosin comparisons. Both delays are significantly different from zero (p-values of 0.031 and 0.0039 for RhoA-myosin and myosin-actin delays, respectively; one-sample Wilcoxon signed-rank test). Scale bars: 10 µm (**A**); 5 µm (**D**,**E)**.

**Supplemental Figure S4:**
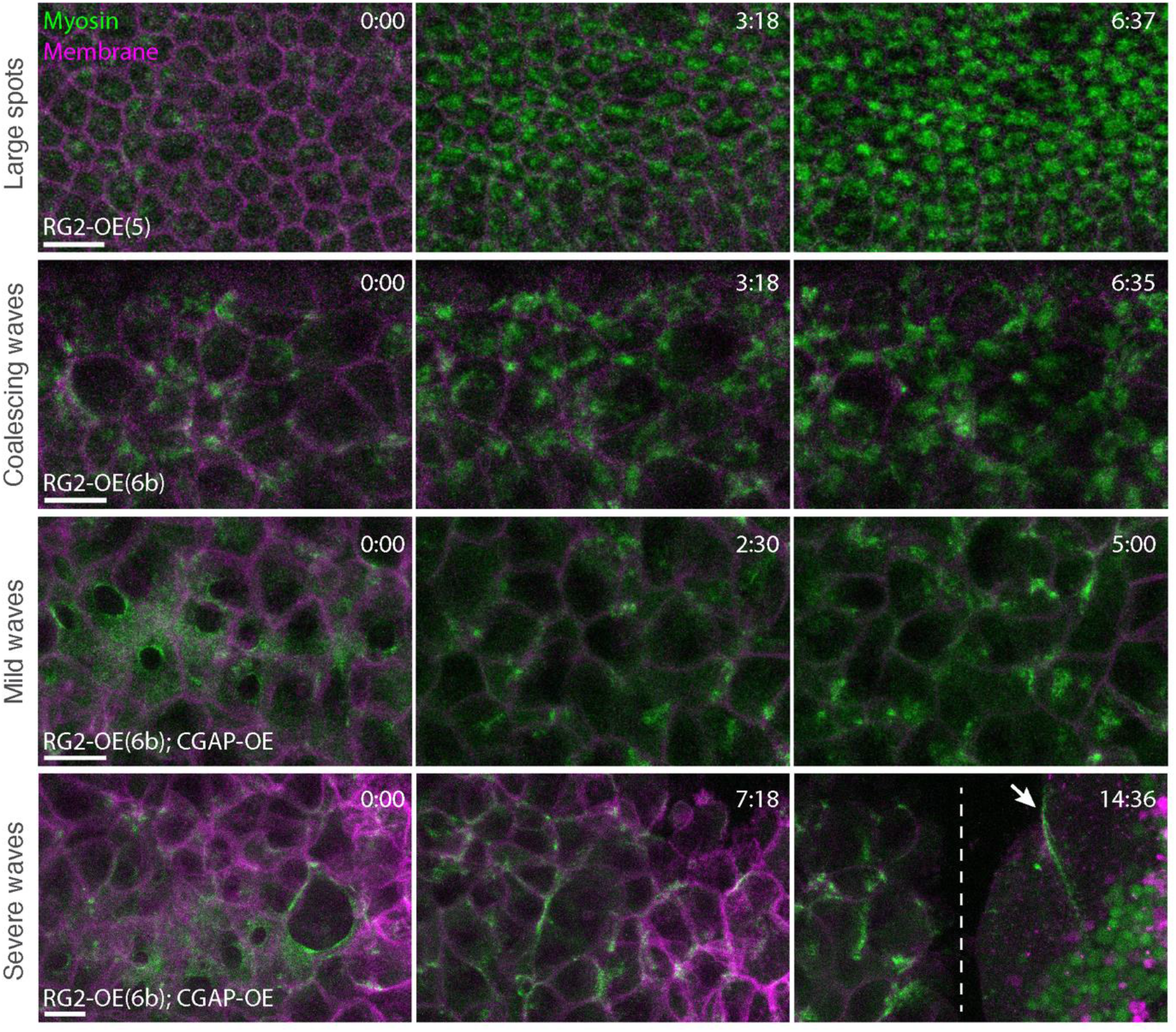
Phenotype classes observed in RhoGEF2-OE and double-OE embryos. MIPs of myosin (green) and membranes (magenta) at three time points for each of the four phenotype classes scored in Figure 5B,C. **A**. Accumulation of abnormally large medioapical spots (‘large spots’). **B**. Waves that form early before transitioning into stable spots and/or a network (‘coalescing waves’). **C.** Waves that appear in some cells after cellularization finishes in part of the embryo (‘mild waves’). These cells are usually pulled towards one pole of the embryo due to cortical instability from cellularization failure at other locations. **D**. Waves that appear in embryos in which incipient cells break down before cellularization is complete in any part of the embryo (‘severe waves’). Third panel shows a large-scale wave deforming the open cortex (arrow) next to the border between open cortex and partially-formed cells (dashed line). Scale bars: 10 µm. All timestamps are min:sec.

**Supplemental Figure S5:**
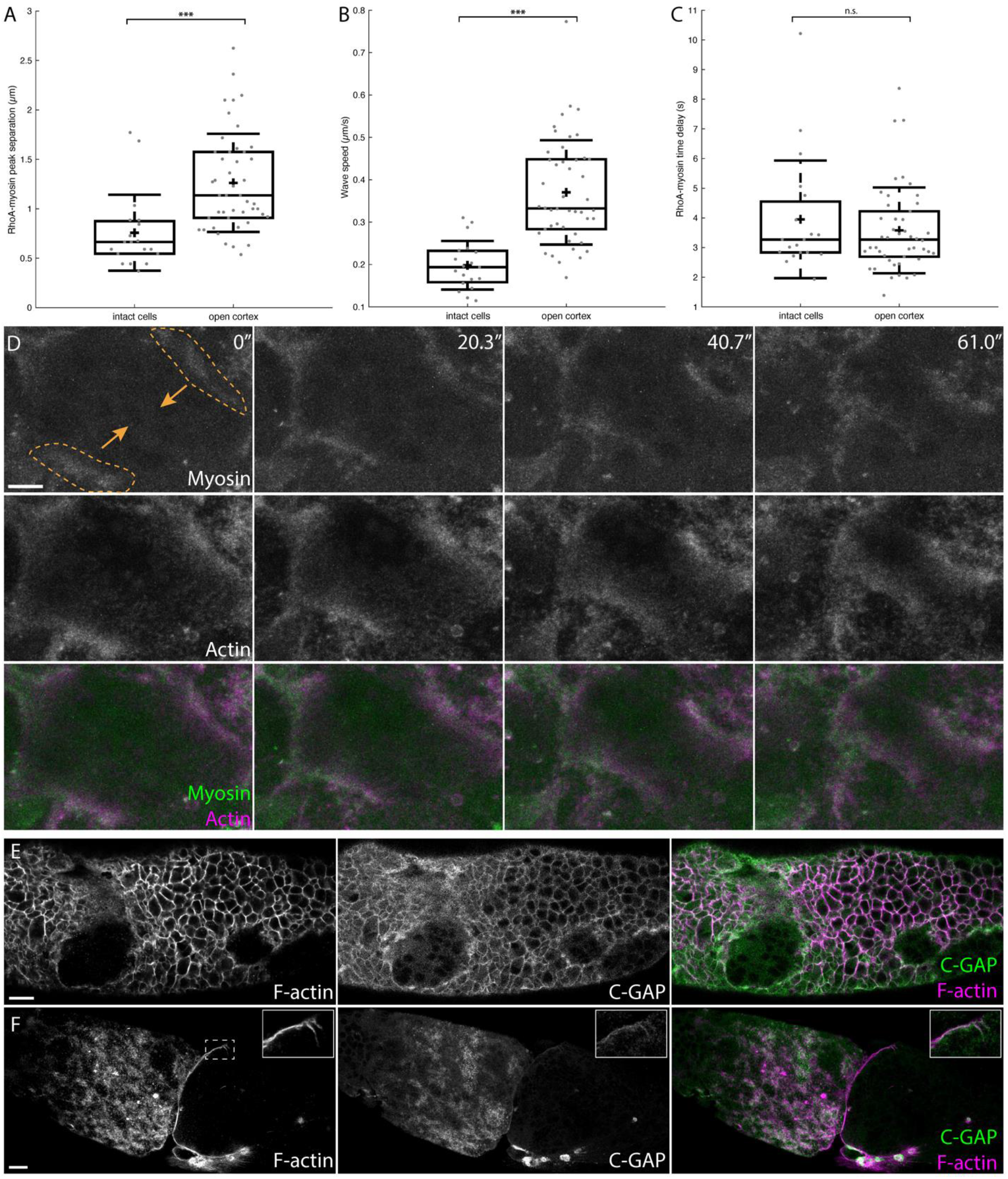
Waves in intact cells and those in open cortex vary in speed but not RhoA-myosin delay, and C-GAP localization is similar to that of F-actin. **A.** Spatial separation between peaks of the RhoA and myosin wavefronts in intact cells and in the open cortex after cellularization fails. **B**. Wave speed for the same two classes of waves. **C**. RhoA-to-myosin delay time for the two classes of waves. All data for **A**-**C** are from double-OE(6b) embryos; the same waves and embryos are used for all three plots. n = 19 waves from 4 embryos for intact cells and 46 waves from 6 embryos for open cortex. Using the Mann-Whitney U test and individual waves as samples, p ∼ 2.9 x 10^−5^, 1.2 x 10^−10^, and 0.72 for **A**, **B**, and **C**, respectively. Box plot components are as in Figure 3C, but without colored points for embryo means. **D.** MIPs of myosin (top), F-actin (middle), and merged channels (bottom) at four time points in a double-OE(6b) embryo. F-actin leads myosin slightly in the pair of converging waves outlined in orange. **E**. F-actin and C-GAP signals from a fixed embryo in which tears were just beginning to form in the epithelial layer. **F**. The same signals from an embryo in which the cells had almost entirely broken down, leaving an open cortex to the right of the image. Scale bars: 10 µm (**D**); 20 µm (**E**,**F**).

The onset of actomyosin waves was variable, often beginning in syncytial embryos and/or causing defects in cellularization, in contrast to the normal medioapical pulses, which began only after cellularization was complete. In the most severe cases, only observed in double-OE embryos, incipient cells tore apart before fully forming and left a continuous cortex spanning much of the embryo’s surface (referred to here as ‘post-breakdown embryos’). As this breakdown occurred, waves organized into large, repeating propagating fronts, which annihilated upon collision (Video S6). We quantified the frequency with which different severities of phenotypes occurred in RhoGEF2-OE or double-OE embryos (Figure 5B,C with different classes of phenotype shown in Supplemental Figure 4). While there was some overlap in phenotypes observed between RhoGEF2-OE and double-OE embryos, double-OE embryos were unlikely to have myosin pulses and often never completed cellularization.

Because temporal delays between components could be read out as spatial displacements between the wavefronts, actomyosin waves in the post-breakdown embryo were a convenient system for investigating the order of protein recruitment. In addition, this system allowed us to assess the excitable behavior of waves induced by RhoGEF2 and C-GAP double-OE, because we could observe wave collisions, spiral waves, and other behaviors that are indicative of excitable behavior. Double-OE(6b) embryos (containing the stronger of the two RhoGEF2-OE insertions) more often displayed cellularization failure and traveling waves than double-OE(5) embryos, so we focused on this genotype. Imaging double-OE(6b) embryos expressing both myosin and anillinRBD following cellularization failure showed both proteins organized into wavefronts, with RhoA preceding myosin (Figure 5D; Video S6). By measuring the wave speed and the distance between the two peaks, we estimated the delay time between RhoA and myosin to be 3.6 +/− 1.0 s, with a wave speed of 0.35 +/− 0.06 µm/s (mean +/− standard deviation; Figure 5F). This result is consistent with RhoA activation inducing actomyosin assembly and with a prior study that found a delay of approximately 10 seconds between RhoGEF2 and myosin pulses in intact mesoderm cells^14^. Additionally, the wave speed measured here is similar to measurements of ∼0.3 µm/s for the wave speed in nurse cells^13^. Although waves in intact cells in less severely-impacted embryos traveled at a lower speed than those in the cortex of post-breakdown embryos, the time delay between RhoA and myosin was not significantly different between the two types of waves (Supplemental Figure 5A-C).

In addition to its requirement in myosin-based contractile force generation, F-actin has also been implicated as a source of negative feedback in other systems displaying cortical excitability^10,15,16^. To better understand the relation of F-actin to myosin in our system, we imaged double-OE(6b) embryos expressing the labeled actin-binding domain of Utrophin (UtrABD) alongside myosin and repeated the analysis above. Interestingly, while on average the myosin signal led that of F-actin by 1.1 +/− 0.6 s (mean +/− s.d.), with wave speeds of 0.42 +/− 0.1 µm/s; Figure 5E-G; Video S6), in a minority of waves, the order was reversed (Supplemental Figure 5D). In fixed double-OE(6b) embryos, C-GAP and F-actin displayed similar patterns both in intact cells and in the open cortex of post-breakdown embryos (Supplemental Figure 5E,F), supporting prior studies and models in which F-actin recruits the GAP to form a negative feedback loop^10,15–17^. Overall, our data suggest that RhoGEF2 and C-GAP can induce excitable actomyosin behavior, similar to Ect2 and RGA-3/4 in *C. elegans*, *Xenopus*, and starfish and to Ect2 and RhoGAP15B in nurse cells.

## Discussion

We discovered that increasing availability of both members of a RhoGEF/RhoGAP pair can induce cortical actomyosin waves in different *Drosophila* cell types. We identified Ect2/Pebble as a RhoGEF involved in nurse cell actomyosin waves during oogenesis, and we showed that Ect2 and RhoGAP15B are released from the nucleus during development in the same order in which cells transition from uniform to wave-like contractility. Similarly, in the early embryo, co-overexpressing a different regulatory pair (RhoGEF2 and C-GAP) leads to waves in epithelial cells and in the open cortex following cellularization failure (Figure 6). By analyzing waves induced in the post-breakdown embryo and comparing the localization of different members of the RhoA pathway, we suggest these waves are similar to pulses and waves in other systems controlled by a different GEF-GAP pair (Ect2 and RGA-3/4). This similarity suggests the RhoA signaling network’s structure and feedback loops give rise to similar regulatory logic for RhoA-mediated excitability across different cell types, GEF/GAP pairs, and developmental contexts.

**Figure 6:**
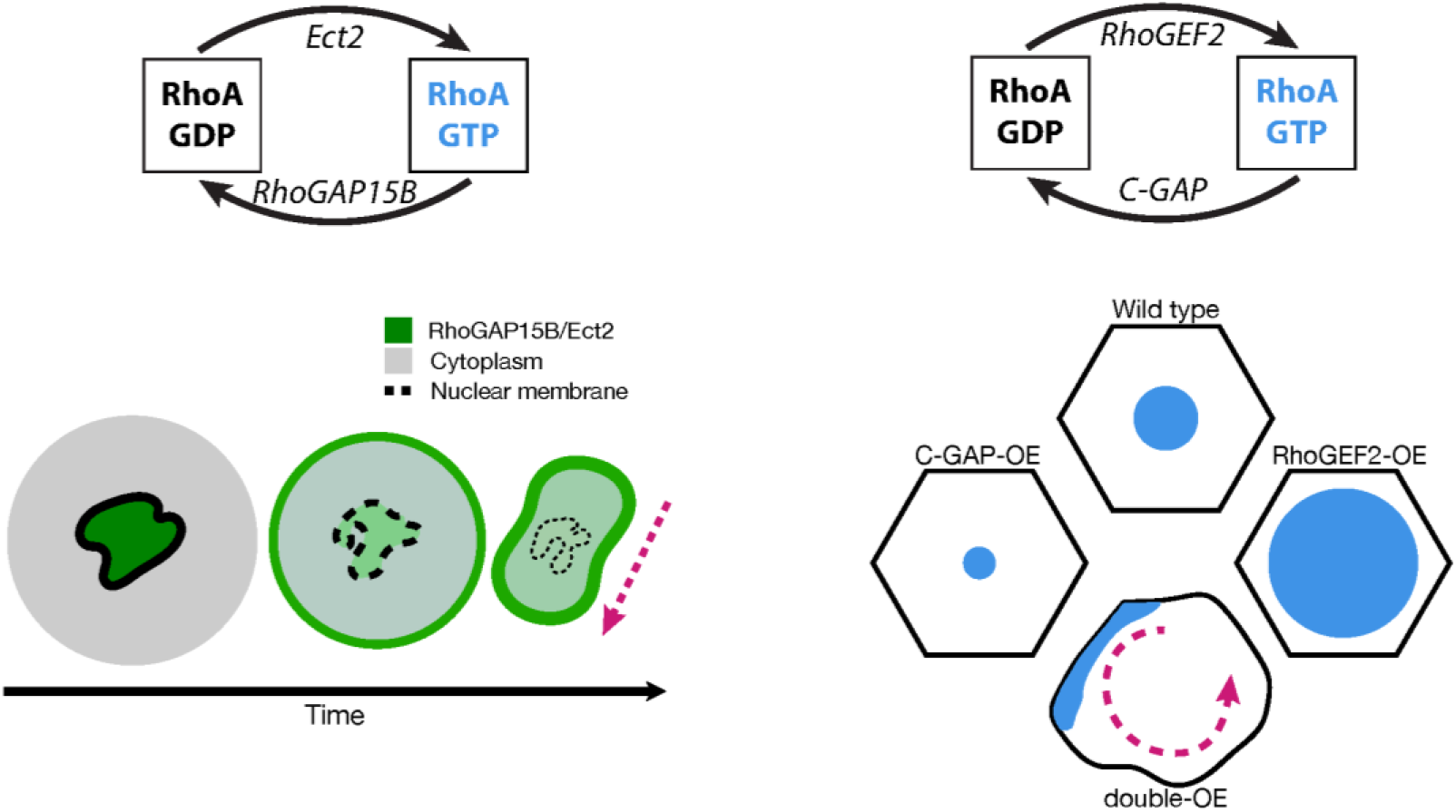
Increase in GAP and GEF levels can induce waves in the actomyosin cortex. (Left) In nurse cells during dumping, Ect2 and RhoGAP15B form a regulatory pair. Both proteins exit the nucleus as dumping progresses, associated with a switch in actomyosin behavior from uniform to wave-like. (Right) In mesoderm cells during ventral furrow formation, RhoGEF2 and C-GAP regulate RhoA. Overexpressing either regulator affects the size of actomyosin pulses, while overexpressing both together converts pulses into waves.

### Precisely regulated cortical excitability is required for development

Regulation of actomyosin contractility through RhoGAP and RhoGEF activity is critical in many developmental contexts^6,41–48^, suggesting GAP/GEF localization changes might drive changes in contractility. Indeed, in nurse cells, although much of cytoplasmic transfer occurs prior to wave onset, the emergence of contractile waves is required for complete cytoplasmic transfer, with perturbations of RhoGAP15B or Ect2 disrupting the wave-mediated phase of transport.

Although our correlative data do not determine whether release of regulators from the nucleus is necessary for wave formation, prior studies of Ect2 suggest a causal role is possible: Ect2 is required both for mitotic rounding and contraction of the cytokinetic ring^35,36,49^. In this context, Ect2 is released from the nucleus shortly before nuclear envelope breakdown, whereupon it promotes myosin wave behavior across the cortex, presumably through the reported wave-like Rok recruitment^13^. Nurse cells are post-mitotic by dumping onset; instead, they begin to undergo a form of programmed cell death mediated by the surrounding somatic cells^50^. In particular, nuclear envelope integrity is compromised, and many components, including PCNA and β-galactosidase attached to a nuclear localization sequence, have been shown to exit the nucleus at this point^13,51,52^ (Figure 3A,B; Supplemental Figure 2A). Thus, cell death-associated loss of nuclear envelope integrity likely releases both GEF and GAP into the cytoplasm, possibly leading to a change in myosin behavior in a process similar to that seen during mitosis.

Speeding or delaying release of Ect2 and RhoGAP15B from the nucleus relative to cell size decrease, either by perturbing the nurse cells themselves or by interfering with the somatic cells responsible for their breakdown, would be an interesting next step to directly test whether nuclear release of these regulators is required for wave onset. However, we showed that decreasing overall levels of either Ect2 or RhoGAP15B disrupts wave onset, suggesting that RhoA signaling output is sensitive to the cytoplasmic levels of these proteins.

In ventral furrow formation, actomyosin pulses present a different excitable behavior that is also required for development but regulated by a different RhoGAP/RhoGEF pair. In GEF/GAP overexpression or knockdown embryos, the furrow is either blocked or formed with abnormal size or shape^14,24^, and when both regulators are overexpressed and waves appear, furrowing is blocked entirely (Video S5). Given the ability of mesoderm cells to generate traveling waves, why are medioapical spots prevalent in the mesoderm rather than waves or some other behavior involving movement of myosin? One possibility is a difference in the activity level of RhoA itself: there could be less RhoA or a higher ratio of GAP:GEF in the mesoderm cells. However, neither injection of constitutively active RhoA nor knockdown of C-GAP leads to waves^14^, and overexpression of RhoGEF2 alone typically leads to larger pulses rather than waves^24^ (Figures 4A and 5B). An alternative hypothesis is that confinement of RhoA, myosin, and/or other components of the RhoA pathway limits their movement across the apical cell cortex and prevents advection of components, leading to a pulse rather than a traveling wave^53^. Nurse cells and other common model systems for RhoA waves and pulsatile contractions are larger, typically lack epithelial polarity, and have few, if any, cell neighbors. In contrast, the epithelial cells forming the ventral furrow have well-defined cell-cell junctions and many neighbors. Consistent with this hypothesis, knockdown of alpha-catenin to disrupt adherens junctions in the mesoderm leads to a contractile instability in which regions of high myosin draw in nearby myosin and spot positioning is less consistent^54^. Although disrupting adherens junctions does not induce waves, an interesting future direction would be to test whether wave formation in adhesion mutants is more sensitive to RhoGEF and/or RhoGAP overexpression. Furthermore, double-OE embryos in which some cells form before cellularization fails often contain a few rounded cells with traveling waves that heavily deform their shapes, similar to nurse cells (Video S6, final portion), suggesting that removal of cell-cell junctions might ease the formation of traveling waves. Finally, differences in levels of downstream effectors could also affect whether a system displays waves or pulses: recent work in *C. elegans* has shown depletion of formin and anillin can convert isolated pulses into oscillatory pulses or noisy, wave-like behavior^55^.

### The cell cortex can transition between different excitable behaviors upon changes to GEF and GAP levels

The transition from isotropic to wave-like myosin in nurse cells is progressive rather than switch-like, with myosin initially organized into small, transient concentrations, reminiscent of pulses, before forming waves that become more intense over time (Supplemental Figure 2B; Video S3). Progression could be driven by a gradual increase in levels of RhoGAP15B and Ect2 as they are released from the nucleus (Figure 3A,B; Video S2), consistent with modeling that suggests changing levels of RhoA regulators or feedback strengths can alter the type of behavior observed^53,56^. The exact mechanism by which higher levels of both GEF and GAP induce waves is unclear. However, previous studies suggest GAP activity increases RhoA flux while GEF activity increases RhoA activity^57,58^, with an increase in both potentially placing the cortex in a state that favors waves.

In yeast cells, polarity sites containing Cdc42-GTP, a Cdc42-GEF, and other proteins have been shown to oscillate and compete with one another until a stable site forms^59^, while in starfish blastomere division, transient waves of RhoA and F-actin appear across the cortex before disappearing as a stable cytokinetic furrow appears^16^. In both cases, transitions between oscillatory or wave-like behavior and stable structures occur, though with different GTPases and in different contexts. A mathematical model for the transition from waves to furrow in the starfish suggests that increasing levels of the GEF (Ect2) at the furrow allows it to outcompete the waves, eventually resulting in a single static structure^56^, providing an example in which change in GEF availability drives transitions in cortical behavior. It is unclear whether a change in RhoGEF2 and/or C-GAP levels over time in wild-type mesoderm cells drives the transition from oscillatory myosin spots to increasing myosin levels^39^, or if changes in endocytosis and/or a physical scaffold like the spectrin cytoskeleton are needed^60,61^.

Although post-breakdown double-OE embryos did not transition between qualitatively different types of contractility (e.g. pulses to waves), they still displayed a progression in the exact pattern of wave-like behavior. Waves at earlier times were often smaller and tended to annihilate quickly. As time progressed, large planar waves frequently emerged and organized into repeating trains of wavefronts, occasionally with transient spiral-wave patterns (Video S6). This progression is similar to that observed in cAMP signaling in fields of aggregating *Dictyostelium* cells, in which short-lived circular waves emerge from spontaneous excitation, with some regions eventually dominating and leading to highly-ordered waves in the cell field^62^. The similarity between the open embryo cortex and cAMP signaling waves further points to general features of excitable systems rather than patterns specific to cell types or specific proteins.

### Other systems displaying cortical excitability suggest potential feedback mechanisms

The origin of the feedback loops underlying excitability in the nurse cells and early embryo are still an area of future interest. Excitability often results from fast positive feedback coupled to delayed negative feedback^18^, and prior studies have suggested several possible mechanisms for such feedback loops in the context of cortical excitability. Active RhoA could feed back positively on itself, as suggested in *C. elegans* and MCF-7 cells^10,11^, possibly through interaction with a RhoGEF. Interestingly, Ect2 and RhoGEF2 have different mechanisms of activation. Ect2 is regulated during cytokinesis by the centralspindlin complex^63–65^, while RhoGEF2 is activated by Gα_12/13_ and a transmembrane protein T48^66^. Thus, this regulatory logic appears to transcend structural and regulatory differences of individual RhoGEF proteins. Alternatively, concentration by advection of components into regions of higher contractility could cause a mechanical form of positive feedback, as suggested in the *Drosophila* germband^8^. While no clear advection of myosin in mesoderm cells was observed, we occasionally observed flows of myosin or F-actin in nurse cells and in the post-breakdown embryo.

In starfish, frogs, and *C. elegans*, inhibition of F-actin has been shown to either disrupt localization of the RhoGAP RGA-3/4 or lead to an increase in RhoA wave amplitude, suggesting F-actin could mediate negative feedback by recruitment of a GAP^10,15,16^. Experiments combining *Xenopus* oocyte extract with a reconstituted artificial actomyosin cortex showed appearance of both waves and oscillations of RhoA and F-actin, with F-actin following RhoA by 15-25 seconds. Combined with observations that waves first appeared in regions of low F-actin, these experiments further support an inhibitory role of F-actin^67^. Indeed, F-actin recruits C-GAP in the *Drosophila* salivary gland^17^, and F-actin and C-GAP appear in similar locations at the waves and at lateral cell membranes in fixed double-OE(6b) embryos (Supplemental Figure 5E,F), suggesting this form of delayed negative feedback could be important in the early embryo. Activated Cdc42 was also seen to form traveling waves during bud polarization in yeast, with wave mobility reduced upon disruption of F-actin or reduction in Cdc42-GAPs^68^. A similar regulatory logic was also proposed for actin waves in migrating cells^69^, suggesting the requirement of both GEF and GAP in traveling waves might extend to other GTPases and organisms as well. Both RhoGAP15B and C-GAP are structurally different from each other and from the RGA-3 and RGA-4 paralogues, with both C-GAP and RGA-3/4’s only known structural domain being the GAP domain^70,71^. Thus, it is not clear what is leading to the delayed recruitment of these proteins and how the different proteins function analogously. Directly investigating whether C-GAP or RhoGAP15B bind F-actin and testing whether RhoA feeds back onto itself would be important future steps to clarify the feedback loops at work in these systems.

## Acknowledgments

We would like to thank the fruit fly community for providing reagents (especially Stefano Di Talia and Eric Wieschaus for PCNA fly lines), as well as FlyBase, the Bloomington Drosophila Stock Center, and the Transgenic RNAi Project at Harvard Medical School. We thank BestGene, Inc. for help with generation of the RhoGAP15B CRISPR mutant, and members of the Martin Lab and Jasmin Imran Alsous for helpful discussions and feedback on the manuscript. This work was supported by the National Institute of General Medical Sciences through grants R01GM125646 and R35GM144115 to ACM.

## Author contributions

J.A.J., M.D.-L., and A.C.M. designed the study. J.A.J. and M.D.-L. performed experiments. J.A.J. and K.A.K. analyzed data with input from M.D.-L. and A.C.M.. J.A.J. and A.C.M. wrote the manuscript with inputs from all authors.

## Materials and Methods

### Fly stocks and crosses

All fly lines used in this study are listed in Supplementary Table 1 along with the crosses used for each figure. For crosses involving RNAi or overexpression, males of the Gal4 driver line were crossed to females containing the knockdown/overexpression construct. For crosses used for egg chamber experiments, ovaries were dissected from F1 females. For embryo experiments, F2 embryos were collected and imaged. Unless otherwise noted, flies were raised at room temperature (approximately 22°C) for egg chamber experiments and 25°C for embryo experiments. For Figure 5B, some F1 flies for the RhoGEF2-OE(5) embryos were raised at 25°C, and some F1 flies for the double-OE(6b) embryos were raised at 18°C or 25°C.

To generate the RhoGAP15B allele missing the RhoGAP domain (*RhoGAP15BΔGAP*), we used CRISPR-Cas9 as previously described^24,72^ to generate a pair of deletions spanning ∼5600 base pairs, including the entire RhoGAP domain but none of the sequence of *CG13000*, a small gene overlapping but in the opposite orientation to *RhoGAP15B*. Two gRNA sequences, GACACCAGGCAAAGCTGACGAGG and AAATTGGTCCTATTGGCCGGAGG (identified using the Find CRISPR tool from Harvard Medical School: https://www.flyrnai.org/crispr/), were cloned into the pCFD5 vector. Vector construction was carried out by VectorBuilder (the vector ID VB210122-1208bzj can be used to find more information on the vector). The plasmid was then injected into y,w; nos-Cas9 flies by BestGene Inc. (Chino Hills, California), with surviving adults crossed to Dr/TM3 flies. Successful deletion was screened for using PCR; the forward primer sequence was CCCAGCAACAATATCAACC, and the reverse primer sequences were GCATCTCAACGGGACGTA and ACTCTGCCTGTGGATGGT for the cut and uncut sequences, respectively.

Interestingly, while RNAi-mediated knockdown of RhoGAP15B in the germarium using the maternal triple driver Gal4 line^73^ entirely prevented formation of viable egg chambers (not shown), flies homozygous for the *RhoGAP15BΔGAP* allele are viable. The exact reason for this viability is not clear, though it’s possible the lack of RhoGAP15B’s RhoGAP activity is compensated for by other proteins, or that some region of RhoGAP15B still present in the RhoGAP15BΔGAP protein plays a role in the germarium. Regardless, we focused in this study on *RhoGAP15BΔGAP* heterozygotes as an alternative method to RNAi for depleting RhoGAP15B’s RhoGAP activity.

### Microscopy

Images were acquired on a Zeiss LSM710 inverted point-scanning confocal microscope, using a 40x/1.2 NA C-Apochromat water objective (except for zoomed-out images of egg chambers in Supplementary Figure 1E,F, which were acquired using a Plan-Apochromat 25x/0.8 water objective). Lasers used include a 488-nm argon ion, a 561-nm diode, and 594- or 633-nm HeNe lasers. Pinhole sizes ranged from 1 to 3 Airy units (usually 1-2 Airy units for embryo images and 3 for egg chamber movies), with z-spacings ranging from 0.5-1.5 µm for embryo and 1-3 µm for egg chamber images.

### Egg chamber culturing and imaging

Egg chamber dissections and *ex vivo* culturing were performed following an established protocol^74^. Prior to dissection, flies of 2-4 days of age were transferred to a new vial containing a layer of dry yeast. Ovaries were removed from ∼3 flies into Schneider’s *Drosophila* medium (Gibco, #21720001); individual egg chambers were then separated and transferred to glass-bottomed dishes (MatTek, #P35G-1.5-14-C) in 200 µL of Schneider’s medium. If used, CellMask Deep Red Plasma Membrane stain (Invitrogen, #C10046) was added at 1:1000 dilution.

### Embryo mounting and imaging

Embryos were collected from apple-juice agar plates covered by plastic cups. Embryos were staged in Halocarbon 27 oil (Genesee Scientific, #59-134) and those of the relevant stage were dechorionated in 50% bleach solution, rinsed twice in water, then mounted on a glass slide covered in glue (Scotch tape resuspended in heptane) inside a channel created between two #1.5 coverslips used as spacers. Embryos were oriented with the ventral side facing away from the glue, then the channel was covered with a #1 coverslip. The chamber was then filled with Halocarbon 27 oil, after which embryos were imaged.

### Fixation and immunofluorescence

Embryos were dechorionated in 50% bleach then fixed in 4% paraformaldehyde (Electron Microscopy Sciences, #15714) in heptane and phosphate buffer for 60 minutes. The vitelline membrane was removed by hand using a needle and embryos were stained with Alexa568-conjugated phalloidin (Invitrogen, #A12380) overnight at 4°C. Embryos were incubated for 2-2.5 hours at room temperature with a rat anti-HA antibody (Roche, #11867423001) at a 1:1000 dilution, then for 1.5 hours at room temperature with goat anti-rat Alexa647 antibody (Invitrogen, #A21247) at a 1:500 dilution, after which embryos were mounted in Aqua-Poly/Mount (Polysciences, Inc., #18606-100) and the slide allowed to cure for at least four hours.

### RhoGEF RNAi screen

The screen for RhoGEFs that play a role in actomyosin waves in nurse cells was performed in two stages. First, each of the 26 *Drosophila* RhoGEFs was independently knocked down by short-hairpin RNA interference, accomplished by crossing mat67>Gal4; mat15>Gal4 driver males to UAS-[RhoGEF-RNAi] females. F1 flies were collected in apple-juice plate cages and maintained at 27°C. Each day for three days, embryos on the plates were observed by eye; lines for which short, misshapen embryos were seen on more than one day were recorded. This round of the screen was repeated, and lines recorded in both rounds were set up a third time. 100-300 embryos per day for three days were counted from this round of the screen (Figure 2A) to determine frequency of the short-embryo phenotype. Because the phenotype had extremely low penetrance and 85-90% of embryos were normal length even among known positive controls, each of these lines was then imaged live (using a mat67>Gal4; sqh::GFP driver) to determine whether waves in nurse cells were perturbed. Of these, Ect2-RNAi gave the most consistently unusual waves; of note, CG30440-RNAi egg chambers were already smaller than wild-type egg chambers prior to dumping onset, suggesting any potential issues occurred an earlier developmental stage. Images in Figure 2B were acquired with the camera of an iPhone aligned with one eyepiece of a Stemi-2000 stereo microscope (Zeiss).

### Image analysis and quantification

Image processing and figure preparation were performed using FIJI^75^ and the Adobe Illustrator 2023 software. Quantification was performed using either FIJI or Matlab (R2021a) using custom-written scripts.

#### RhoGAP15B / Ect2 relocalization

Relocalization of RhoGAP15B and Ect2 from the nucleus to the cytoplasm/cortex was determined by measuring the mean intensity of either Ect2::EGFP or RhoGAP15B::sfGFP in manually-drawn regions of interest (ROIs) in a single slice 9-12 µm below the top surface of the nucleus. ROIs were drawn using FIJI’s Polygon Selection tool for the nuclei and the Segmented Line tool with a line width of 3 pixels for the cortex. ‘Early’ timepoints were taken from nurse cells prior to visible volume decrease (approximately at dumping onset), while ‘late’ timepoints were taken around the start of wave onset or when the nucleus was no longer clearly distinguishable from the cytoplasm by eye. The mean intensity in each ROI was calculated and the ratio taken for each cell at each timepoint.

#### PCNA leakage and wave onset determination

To determine the order of nuclear breakdown, PCNA::GFP was used as a proxy for Ect2 as it was less perturbative to the egg chambers. ‘Nuclear exit’ was defined as the time at which the cytoplasm-to-nucleus intensity ratio 9-12 µm below the nucleus surface exceeded 7.5%, which roughly corresponded to the median ratio at which PCNA was determined by eye to be ‘in the cytoplasm’ for several cells. ‘Wave onset’ was determined by eye using the CellMask stain to visualize fluctuations in membrane position associated with traveling actomyosin waves. Since there is variability in the rate at which nurse cells transfer their contents in different egg chambers, as well as differences in the number of visible cells in different movies, times of nuclear exit and wave onset were ranked, and the ranks were pooled into a heatmap (Figure 3E); rank-ordering measurements also reduces potential error stemming from estimating wave onset by eye.

#### RhoGEF2-OE and double-OE lines

The two RhoGEF2 overexpression fly lines used in this study contain the same insertion (T7-tagged RhoGEF2 under UASp control) on the second (RG2-OE(6b)) or third (RG2-OE(5)) chromosome (FlyBase reference FBrf0191631). Double-OE(5) and double-OE(6b) fly lines were made using each of the two insertions. Flies containing the RG2-OE(6b) insertion tended to have more severe phenotypes than their RG2-OE(5) counterparts: waves appeared more often upon expression of RhoGEF2 alone when using that line, and cellularization failed earlier and more often when RhoGEF2 was co-overexpressed alongside C-GAP in the double-OE(6b) line than the double-OE(5) line (Figure 5B,C). Unless otherwise mentioned, all double-OE embryos shown in figures and videos contain the RG2-OE(6b) insertion.

#### Embryo overexpression phenotype scoring

Scoring of phenotypes in RhoGEF2 and/or C-GAP overexpressing embryos was performed manually according to the classes defined in the main text and in Supplemental Figure 4). Phenotype scores were recorded, then imported into Matlab to generate bar plots (Figure 5B,C).

#### Wave speed and delays between wave components

Waves with clear signal in both channels were identified manually and an ROI along their path for 3-5 frames drawn by hand. Line widths of ∼10-80 pixels, depending on wave width perpendicular to the propagation direction, were chosen to minimize noise-induced fluctuation in the signal while avoiding capturing signal from other nearby waves. Wave speed was calculated by measuring the position of the wave front at the start and end of the ROI and dividing by the elapsed time. Plots of the intensity of both proteins along the ROI were generated using the ‘RGB_Profiler’ plugin (https://imagej.nih.gov/ij/plugins/rgb-profiler.html), from which the lag distance between peaks in the two channels was measured by hand. Delay time was calculated as the ratio of lag distance to wave speed.

#### Myosin area fraction measurement

To calculate the myosin coverage fraction in different genotypes, maximum intensity projections were made through approximately 10 µm from the ventral surface of each embryo. A single timepoint was extracted from each image sequence, corresponding to roughly 7.5 minutes after the first myosin pulse appeared. Due to variability in constriction rate among genotypes, timepoints ranged from 6 to 9 minutes after the first visible pulse, chosen to standardize the degree of constriction between embryos. Double-OE embryos never began to form a furrow, with myosin first appearing during cellularization, so instead timepoints were chosen shortly before cell breakdown occurred.

The ‘Pixel classification + Object classification’ workflow of Ilastik was used to create a mask of myosin spots for embryos during gastrulation. Images were blinded before program training began. The program was trained on 3 zoomed-in images from each genotype and an additional 3 pairs of zoomed-out images from different genotypes. Sigmas of 1,1.6,3.5 and 5 were selected. During pixel classification training several spots of different sizes and shapes were selected for foreground and several undesired features were selected as background. For thresholding the image the settings of hysteresis, 0 smooth, and threshold of core .3, final .35 as well as a broad size filter (min: 2, max: 1000000) to accept any size spot were selected. For object classification, multiple correct and incorrect objects of various shapes and sizes were selected for each image.

Some data (1 of 22 RhoGEF2-knockdown embryos and 2 of 13 double-OE embryos) resulted in poor outputs where background was clearly interpreted as foreground, nearly the entire embryo was selected as foreground, or nearly all myosin spots clearly identifiable by eye were not identified by Ilastik. These images were excluded from analysis.

The output masks were then used to calculate fractional coverage of the mesoderm cells. The membrane channel of each image was used to select ROIs of approximately 4 cells wide covering the mesoderm cells or to select regions 4 cells wide in double-OE embryos. These regions were used on the output predictions to calculate the fraction of pixels inside the ROI corresponding to foreground (the myosin coverage fraction).

#### Statistical tests

All statistical tests were performed using the ranksum() (for Mann-Whitney U test) and signrank() (for Wilcoxon signed-rank test) functions in Matlab. Samples used in the tests were the mean values per egg chamber or embryo, except for the tests in Supplemental Figure 5A-C. For these measurements, individual embryos showed a mix of wave types and the relevant question was whether waves in intact cells and in open cortex had different characteristics, so the waves, which were assumed to be independent, were used as samples.

## Video captions

**Video S1: Effects of RhoGAP15B and Ect2 perturbation on actomyosin waves in nurse cells**. Maximum-intensity projections of Sqh::GFP for egg chambers of the following six genotypes: wild-type, *RhoGAP15BΔRhoGAP/+*, RhoGAP15B RNAi, *pbl*^3^*/+*, Ect2 RNAi, and Ect2-OE, corresponding to egg chambers used for Figure 2 and Supplemental Figure 1. All RNAi and overexpression constructs are germline-specific. Scale bars: 50 µm; time stamps are hr:min:sec; t=0 is used only for reference and is not standardized between movies. Time steps are different between movies and range from 12 to 30 seconds.

**Video S2: Ect2 and RhoGAP15B exit the nucleus as dumping proceeds**. First movie: MIP of Ect2 under control of the *sqh* promoter during dumping, showing exit from the nucleus in posterior nurse cells before anterior ones. Second movie: a similar pattern is seen with RhoGAP15B. Scale bars: 50 µm; time stamps are hr:min:sec. Time step is 1 minute for the first movie and 2.5 minutes for the second.

**Video S3: Myosin contractions increase in intensity and become more wave-like as dumping proceeds.** Two concatenated time series of the same egg chamber, showing progression in myosin contractility from local spots to waves that propagate across the cell. Scale bars: 50 µm; time stamps are hr:min:sec.

**Video S4: Waves are observed in egg chambers expressing germline-specific Zip::GFP**. MIPs from egg chambers expressing germline-specific Zip::GFP (left) and ubiquitous Sqh::GFP (right), showing waves are inside nurse cells, despite some overlying follicle-cell specific myosin dynamics. All movies are from the same egg chamber; scale bars: 50 µm in the first two movies and 20 µm in the last; timestamps are hr:min:sec. Time step in the last movie is 12 seconds and 20 seconds in the first two.

**Video S5: Myosin behavior in mesoderm cells cells in the genotypes explored in this study**. Myosin pulses appear in wild-type, C-GAP-OE, and RhoGEF2-OE embryos, although pulse size and degree of furrow formation vary with genotype. The two bottom double-OE embryos show relatively mild examples of the traveling wave phenotype, although cells begin to rip apart in the bottom-right movie. Scale bars: 20 µm; time stamps are min:sec. Time steps are different between movies and range from roughly 7 to 18 seconds.

**Video S6: Large traveling waves that annihilate upon collision appear in the cortex remaining after cellularization failure in double-OE embryos**. First movie: myosin signal in a double-OE(5) embryo, with remnant cells to the right. Second movie: zoom-in of myosin waves in a double-OE(6b) embryo, highlighting spiral wave patterns coexisting with planar waves, followed by a whole-embryo view of trains of planar waves. Third movie: myosin and RhoA in a double-OE(6b) embryo, from cellularization failure through emergence of large-scale planar waves, in which RhoA wavefronts can be seen to lead myosin wavefronts. Fourth movie: myosin and F-actin in a double-OE(6b) embryo, with remnant cells to the right. Myosin very slightly leads F-actin in most waves as they gradually become organized into trains of wavefronts. Scale bars: 20 µm; time stamps are hr:min:sec. Time steps vary between movies and range from 9 to 25 seconds.

