## Supplementary Table 1 for "Change in RhoGAP and RhoGEF availability drives transitions in cortical patterning and excitability in *Drosophila*"

| **Fly lines** | | | |
| --- | --- | --- | --- |
| **Fly line number** | **Genotype** | **Target (if RNAi)** | **Source*** |
| 1 | Oregon R |  | Lab stocks |
| 2 | sqh::GFP (chromosome II) |  | Lab stocks |
| 3 | Ubi>GFP::anilRBD/TM3 |  | Lab stocks |
| 4 | w*; P{sqh-pbl-EGFP}30 |  | 76258 |
| 5 | PCNA::GFP (chromosome III) |  | Gif from Dr. Eric Wieschaus |
| 6 | w*; Sp/CyO; PCNA::tagRFP/TM3 |  | Gift from Dr. Stefano Di Talia |
| 7 | RhoGAP15B::sfGFP |  | VDRC 318338 |
| 8 | ubi>Pav::mCherry/TM3 |  | Gift from Dr. Emmanuel Derivery |
| 9 | w*; pbl3/TM3, Sb1 |  | 9358 |
| 10 | RhoGAP15B deletion mutant |  | This study |
| 11 | mat67>GAL4; mat15>GAL4 |  | Lab stocks |
| 12 | mat67>GAL4; sqh::GFP |  | Lab stocks |
| 13 | mat67>GAL4,sqh::mCherry; Dr/TM3 |  | Lab stocks |
| 14 | mat67>GAL4,sqh::GFP/CyO; Gap43::mCherry/TM3 |  | Lab stocks |
| 15 | sqh::GFP/CyO; mat15>GAL4,Gap43::mCherry/TM3 |  | Lab stocks |
| 16 | mat67>GAL4,sqh::GFP/CyO; mat15>GAL4,Gap43::mCherry/TM3 |  | Lab stocks |
| 17 | mat67>GAL4; Ubi>GFP::anillinRBD/TM3 |  | Lab stocks |
| 18 | mat67>GAL4,sqh::mCherry/CyO; Ubi>GFP::anillinRBD/TM3 |  | Lab stocks |
| 19 | mat67>GAL4,UtrophinABD::GFP/CyO; mat15>GAL4,sqh::mCherry/TM3 |  | Lab stocks |
| 20 | tj>Gal4 |  | Lab stocks |
| 21 | UAS-Zipper::GFP |  | Lab stocks |
| 22 | w*; P{UAS-HA-pbl}2m |  | 66161 |
| 23 | y[1] w[*]; P{w[+mC]=UASp-T7.RhoGEF2}5 |  | 9386 |
| 24 | y[1] w[*]; P{w[+mC]=UASp-T7.RhoGEF2}6b |  | 9387 |
| 25 | M{UAS-RhoGAP71E.ORF.3xHA}ZH-86Fb |  | FlyORF (F001171) |
| 26 | P{UAS-T7::RhoGEF2}5, M{UAS-RhoGAP71E.ORF.3xHA}ZH-86Fb |  | Lab stocks |
| 27 | P{UAS-T7::RhoGEF2}6b; M{UAS-RhoGAP71E.ORF.3xHA}ZH-86Fb |  | Lab stocks |
| 28 | y1 sc* v1 sev21; P{TRiP.HMS00412}attP2 | RhoGAP71E | 32417 |
| 29 | y[1] sc[*] v[1] sev[21]; P{y[+t7.7] v[+t1.8]=TRiP.HMS01118}attP2 | RhoGEF2 | 34643 |
| 30 | y1 v1; P{TRiP.HMJ02093}attP40/CyO | RhoGAP15B | 42527 |
| 31 | y[1] sc[*] v[1] sev[21]; P{y[+t7.7] v[+t1.8]=TRiP.GL00344}attP2/TM3, Sb[1] | hts | 35421 |
| 32 | y[1] sc[*] v[1] sev[21]; P{y[+t7.7] v[+t1.8]=TRiP.GL01052}attP2 | Rh3 | 36885 |
| 33 | y[1] sc[*] v[1] sev[21]; P{y[+t7.7] v[+t1.8]=TRiP.GL01092}attP2 | pbl | 36841 |
| 34 | y[1] v[1]; P{y[+t7.7] v[+t1.8]=TRiP.JF02979}attP2 | pbl | 28343 |
| 35 | y[1] v[1]; P{y[+t7.7] v[+t1.8]=TRiP.JF01651}attP2 | CG15611 | 31158 |
| 36 | y[1] v[1]; P{y[+t7.7] v[+t1.8]=TRiP.JF01725}attP2 | CG30440 | 31207 |
| 37 | y[1] sc[*] v[1] sev[21]; P{y[+t7.7] v[+t1.8]=TRiP.GL00482}attP2 | CG30440 | 35635 |
| 38 | y[1] v[1]; P{y[+t7.7] v[+t1.8]=TRiP.JF01659}attP2/TM3, Sb[1] | CG42674 | 31166 |
| 39 | y[1] v[1]; P{y[+t7.7] v[+t1.8]=TRiP.HMJ23766}attP40/CyO | CG43102 | 62371 |
| 40 | y[1] v[1]; P{y[+t7.7] v[+t1.8]=TRiP.JF03182}attP2 | CG43658 | 28754 |
| 41 | y[1] sc[*] v[1] sev[21]; P{y[+t7.7] v[+t1.8]=TRiP.HMS00332}attP2 | CG43658 | 32341 |
| 42 | y[1] sc[*] v[1] sev[21]; P{y[+t7.7] v[+t1.8]=TRiP.HMS01370}attP2/TM3, Sb[1] | CG46491 | 34380 |
| 43 | y[1] v[1]; P{y[+t7.7] v[+t1.8]=TRiP.JF01661}attP2 | Cdep | 31168 |
| 44 | y[1] sc[*] v[1] sev[21]; P{y[+t7.7] v[+t1.8]=TRiP.GL00650}attP2 | cyst | 41578 |
| 45 | y[1] sc[*] v[1] sev[21]; P{y[+t7.7] v[+t1.8]=TRiP.HMS00246}attP2 | Exn | 33373 |
| 46 | y[1] v[1]; P{y[+t7.7] v[+t1.8]=TRiP.HMJ02116}attP40 | GEFmeso | 42545 |
| 47 | y[1] v[1]; P{y[+t7.7] v[+t1.8]=TRiP.GLC01361}attP40 | mbc | 44424 |
| 48 | y[1] sc[*] v[1] sev[21]; P{y[+t7.7] v[+t1.8]=TRiP.HMS00320}attP2 | PsGEF | 33433 |
| 49 | y[1] v[1]; P{y[+t7.7] v[+t1.8]=TRiP.JF01776}attP2 | Pura | 31221 |
| 50 | y[1] sc[*] v[1] sev[21]; P{y[+t7.7] v[+t1.8]=TRiP.HMS00267}attP2/TM3, Sb[1] | RhoGAP1A | 33390 |
| 51 | y[1] v[1]; P{y[+t7.7] v[+t1.8]=TRiP.JF01747}attP2 | RhoGEF2 | 31239 |
| 52 | y[1] v[1]; P{y[+t7.7] v[+t1.8]=TRiP.JF01686}attP2 | RhoGEF4 | 31178 |
| 53 | y[1] v[1]; P{y[+t7.7] v[+t1.8]=TRiP.JF01604}attP2 | RhoGEF64C | 31130 |
| 54 | y[1] sc[*] v[1] sev[21]; P{y[+t7.7] v[+t1.8]=TRiP.HMC06563}attP40 | RhoGEF64C | 77431 |
| 55 | y[1] v[1]; P{y[+t7.7] v[+t1.8]=TRiP.JF01153}attP2 | RhoGEF3 | 31580 |
| 56 | y[1] sc[*] v[1] sev[21]; P{y[+t7.7] v[+t1.8]=TRiP.HMS00741}attP2 | RtGEF | 32947 |
| 57 | y[1] v[1]; P{y[+t7.7] v[+t1.8]=TRiP.JF01795}attP2 | sif | 25789 |
| 58 | y[1] v[1]; P{y[+t7.7] v[+t1.8]=TRiP.JF01680}attP2 | Sos | 31174 |
| 59 | y[1] sc[*] v[1] sev[21]; P{y[+t7.7] v[+t1.8]=TRiP.HMS01356}attP2/TM3, Sb[1] | spg | 34367 |
| 60 | y[1] v[1]; P{y[+t7.7] v[+t1.8]=TRiP.JF02815}attP2 | trio | 27732 |
| 61 | y[1] sc[*] v[1] sev[21]; P{y[+t7.7] v[+t1.8]=TRiP.HMS01979}attP2 | Vav | 39059 |
| 62 | y[1] v[1]; P{y[+t7.7] v[+t1.8]=TRiP.JF02839}attP2 | Zir | 28005 |
| 63 | y[1] v[1]; P{y[+t7.7] v[+t1.8]=TRiP.HMJ21536}attP40 | Ziz | 54817 |
| *Numbers with no other information correspond to Bloomington Drosophila Stock Center stock numbers; VDRC = Vienna Drosophila Resource Center. | | | |

| **Crosses** | | |
| --- | --- | --- |
| **Figure panel(s) or video** | **Male** | **Female** |
| Fig. 2A, 2B, RhoGEF screen | 11 | 30-63 |
| RhoGEF screen live imaging | 12 | 32-34,36,39,40,53 |
| Fig. 2C | 2 or 8 | 8 or 2 |
| Fig. 2D | 12 | 30 |
| Fig. 2E | 12 | 33 |
| Fig. 2F | 12 | 22 |
| Fig. 3A,A',C | 4 | 4 |
| Fig. 3B,B',C | 7 | 7 |
| Fig. 3D,E | 5 | 5 |
| Fig. 4A,B | 15,16 | 25,29,32,28,23 |
| Fig. 4C | 16 | 27 |
| Fig. 5A | 14,16 | 1,27 |
| Fig. 5B,C | 12,15,16 | 23,24,26,27 |
| Fig. 5D,F,G | 18 | 27 |
| Fig. 5E,F,G | 19 | 27 |
| Fig. S1A | 12 | 1 |
| Fig. S1B | 12 | 30 |
| Fig. S1C | 12 | 33 |
| Fig. S1D | 12 | 22 |
| Fig. S1E | 12 | 30 |
| Fig. S1F | 12 | 33 |
| Fig. S1G | 18 | 22 |
| Fig. S1H | 12 | 10 |
| Fig. S1I | 12 | 9 |
| Fig. S2A | 4 or 6 | 6 or 4 |
| Fig. S2B | 2 or 8 | 8 or 2 |
| Fig. S2C-E | 13 | 21 |
| Fig. S3A | 18 | 1 |
| Fig. S3B | 18 | 23 |
| Fig. S3C | 18 | 24 |
| Fig. S3D | 18 | 27 |
| Fig. S3E | 14 | 1 |
| Fig. S3F | 16 | 25 |
| Fig. S3G | 16 | 23 |
| Fig. S3H | 16 | 27 |
| Fig. S4 | 14,16 | 23,24,27 |
| Fig. S5A-C | 18 | 27 |
| Fig. S5D | 19 | 27 |
| Fig. S5E,F | 17 | 27 |
| Video S1 | 2 | 8,9,10,22,33 |
| Video S2 | 13 | 21 |
| Video S3 | 4,7, (4 or 6), (2 or 8) | 4,7, (6 or 4), (8 or 2) |
| Video S4 | 14,16 | 1,23,25,27 |
| Video S5 | 12,18,19 | 26,27 |
